# Single-cell spatial transcriptome reveals pathological features of human hippocampus with sclerosis

**DOI:** 10.1101/2025.01.21.633854

**Authors:** Lifang Wang, Xingjie Li, Yuyang Liu, Bufan Jin, Yanrong Wei, Youzhe He, Yaru Ma, Zihan Wu, Xue Gong, Yanru An, Xueying Kong, Yanan Xing, Ruixi Ye, Quyuan Tao, Aiqing Li, Mingkai Chang, Li Pu, Jinmei Li, Li Pang, Dong Zhou, Cai Song, Shiping Liu, Xun Xu, Longqi Liu, Lei Han, Zhen Hong

**Affiliations:** Department of Neurology, West China Hospital of Sichuan University, Chengdu, Sichuan, China; BGI Research, Hangzhou 310030, China; BGI Research, Shenzhen 518083, China; College of Life Sciences, University of Chinese Academy of Sciences, Beijing 100049, China; Institute of Brain science and Brain-inspired technology of West China Hospital, of Sichuan University, Chengdu, Sichuan, China; Tencent AI Lab, Shenzhen 518057, China; BGI Research, Qingdao 266555, China

## Abstract

More than half of epileptic patients ultimately turned to intractable epilepsy. Mesial temporal lobe epilepsy with hippocampal sclerosis (MTLE-HS), the most common type of intractable epilepsies, whose pathological mechanism remains elusive. Here, using 42 human hippocampal samples from surgical donors of MTLE (32 with and 10 without HS) through single-cell resolution Stereo-seq and histologcial experiments, we revealed spatially pathological changes of gene expression and cell type composition with HS systematically. After precise parcellation of hippocampal subregions and differentially expressed gene (DEG) analysis between each region with and without sclerosis, we found Cornu Ammonis (CA) subregions with higher number of DEGs were vulnerable to sclerosis, especially CA1 and CA3. Within CA1, we found that CA1-superficial and proximal areas were more vulnerable than CA1-deep and distal areas in sclerosis. Meanwhile, after analyzing 350,795 segmented cells from Stereo-seq, we found dramatically increasing density of astrocyte accompanied with significantly decreasing density of excitatory neurons in CA1, especially superficial and proximal CA1, and CA3 in sclerosis. In these two vulnerable subregions, proliferative astrocyte (P_astrocyte) and reactive astrocyte (R_astrocyte) were found to be enriched whereas apoptotic subtype of astrocyte (A_astrocyte), related with apoptotic pathway, was mainly located in alveus, which strengthened cell communication with reactive microglia (R_microglia) in HS, revealing the novel pathological feature in our work. The pseudotime analysis indicated that CA excitatory neurons underwent synaptic impairment, energy dysfunction, aging, and finally losing cell identity until death through autophagy or apoptosis. Besides, we also found a resilient subtype, EX_CA2-4.3, highly expressed extracellular matrix related genes including *PDYN*, and was increasing the interaction of BDNF-NTRK, NFASC-CNTN1 to withstand the damage from sclerosis. Together, our study provided a reference of human hippocampus with and without HS caused by MTLE, and highlighted the potential pathological mechanism on molecular and cellular level of MTLE-HS.

## Introduction

Mesial temporal lobe epilepsy with hippocampal sclerosis (MTLE-HS), the most common type of intractable epilepsies^1–3^, was a ponderous burden for both patients and healthcare system. To prevent fast worsening of the disease condition and/or sudden death occurrence^4^, neurosurgical resection of epileptic lesion, normally temporal lobe and hippocampus, was the last procedure, which aggregated the declarative memory deficits.

In general, HS was related to distinct patterns of hippocampal damage^3,5,6^, including variable extents of astrogliosis and neuronal loss^6,7^. The advent of single-cell/single-nucleus RNA sequencing (sc/snRNA-seq) facilitated the understanding of potential molecular and cellular pathology of epilepsy. SnRNA-seq of temporal lobe from TLE patients showed that the ratio of neurons was decreased whereas the ratio of astrocyte and microglia was increased, they also found a type of reactive astrocyte which played a role in promoting neuronal hyperactivity and disease progression^8^. SnRNA-seq of CA1 neurons from MTLE-HS mouse model was first reported that excitatory neurons located in CA1-superfical layer (CA1-sup) were more vulnerable to HS than CA1-deep layer, and that neurons in CA1-sup of HS undertook neurodegenerative transcriptional trajectories. The expression of death-associated genes and neurodegenerative genes were especially high in CA-sup neurons of HS^9^, suggesting different vulnerability of CA1 subregions in HS mouse model. However, mouse model of HS was quite different from human who suffered seizure spontaneously, and what kind of pathological changes in all hippocampal cells with sclerosis were needed to be systematically explored.

Transcriptomic profile of the lesions for epileptic patients was also much less reported, and mainly focused on the temporal lobe^8^. Francois et al. identified the gene expression profile and regulatory networks for variable intractable epilepsies and suggested that intractable epilepsies shared key potential mechanisms, such as dysfunction of synaptic transmission, synaptic plasticity, immune response, extracellular matrix, and energy derivation. However, these epilepsy-associated transcriptomic change lacked spatial and cellular resolution. As the spatial information for gene expression was crucial for MTLE-HS, Busch et al. tried to dissect the molecular and subregion mechanism of MTLE-HS patients^10^, and found *BDNF* is the pivotal gene in regulatory network in CA3 subregion, however, the number of genes detected was quite limited by using *in situ* hybridization. Since the advent of whole genome based spatial transcriptomics with single-cell resolution (Stereo-seq) in 2022^11^, it is possible and eagerly to study the spatial vulnerability and heterogeneity of MTLE-HS pathology and the spatial cell type composition change systematically.

In this study, Stereo-seq combined with snRNA-seq and histological staining experiments were performed to dissect the spatial distribution changes of gene expression and cell type composition of human hippocampus with sclerosis for the first time. The vulnerability of hippocampal subregions and cell types to HS were systematically investigated. CA1, especially proximal and superficial CA1, and CA3 were the most vulnerable subregions in HS. The density of specific subtypes of astrocyte was increased in CA subregions, especially proximal and superficial CA1. Excitatory neurons undergone neurodegenerative trajectory or resilient trajectory in sclerosis. These findings provided important information for understanding the pathological mechanism in MTLE-HS.

## Results

### Spatially-resolved transcriptome of human hippocampus

To characterize the spatial cellular and transcriptomic differences between human hippocampus with and without sclerosis of MTLE, we collected 42 human hippocampus samples, including 32 samples with sclerosis and 10 samples without sclerosis, from surgical donors with various intractable epilepsies. Whether hippocampus with or without sclerosis was diagnosed under various staining by experienced neurobiologists and pathologists (see **Methods**) (**Fig.1a, Extended data Fig.1, Supplementary Table 1**). These samples have no significant differences in sex (female (54.8 %), *p* = 0.590), seizure types (*p* = 0.335) and surgery side (*p* = 0.878) (**Supplementary Table 2**). Seven qualified samples (including 4 sclerosis (-) and 3 sclerosis (+), RNA integrity index (RIN) > 6) were used for Stereo-seq and snRNA-seq. 10 μm and its neighboring 500 μm thick from the same hippocampal sections were employed for Stereo-seq and snRNA-seq, respectively^11^ (**Fig. 1a, Supplementary Table 3**). The left slices from 7 sequencing samples and other 34 samples were applied for histological staining. After running standard sequencing pipeline, the quality of Stereo-seq data was assessed in the form of bin100 size (100 × 100 DNA nanoballs (DNB) spots, dimension size: 50 µm × 50 µm)^12^. On average, we obtained 2,291 genes/bin100 and 6,692 unique molecular identifiers (UMIs)/bin100 across all hippocampal sections (**Supplementary Table 5**). The hippocampal subregions were parcellated by integration of spatial clustering with BayesSpace algorithm, the nucleus staining, particles of total RNA accumulation, and further adjusted by spatial distribution of cell types^13^ (**Fig. 1b, Extended data Fig. 2a**). Our data revealed a total of eight precisely parcellated anatomical subregions within the hippocampus, including the Cornu Ammonis (CA)1, CA2, CA3, CA4, dentate gyrus (DG), subiculus (Sub) and glial cell regions comprising FAS (fimbria, alveus, and stratum oriens (SO)) and SLRM (stratum lucidum (SL), stratum radiatum (SR), stratum moleculare (SM)) (**Fig. 1b**). We found the gene abundance was various among hippocampal subregions, from lowest subregions with 1,855 genes/bin100 in FAS and SLRM, to highest subregion with 3,919 genes/bin100 in CA2. Besides, the gene abundance of CA subregions in sclerosis were lower than that without sclerosis (**Extended data Fig. 2b)**. After parcellation of subregions in hippocampus, we obtained a series of subregion-specific genes, such as *PDYN* for DG, *MET* for CA1^14^, *SCGN* for CA2, *COX7A1* for CA3-4, *RELN* for SLRM and *BCAS1* for FAS (**Fig. 1c and d, Supplementary Table 6**). We also used two public datasets to verify the accuracy of our subregion parcellation, as expected, our parcellation is consistent with previous reports which parcellated based on histological anatomy (**Extended data Fig. 2c and d**)^14,15^. The expression of subregion specific genes revealed that relatively lower specificity of subregion specific genes in CA1-4 with sclerosis than that without sclerosis (**Fig. 1e-g, Extended data Fig. 2e**), implying pronounced pathological changes in CA region with sclerosis.

**Fig. 1.**
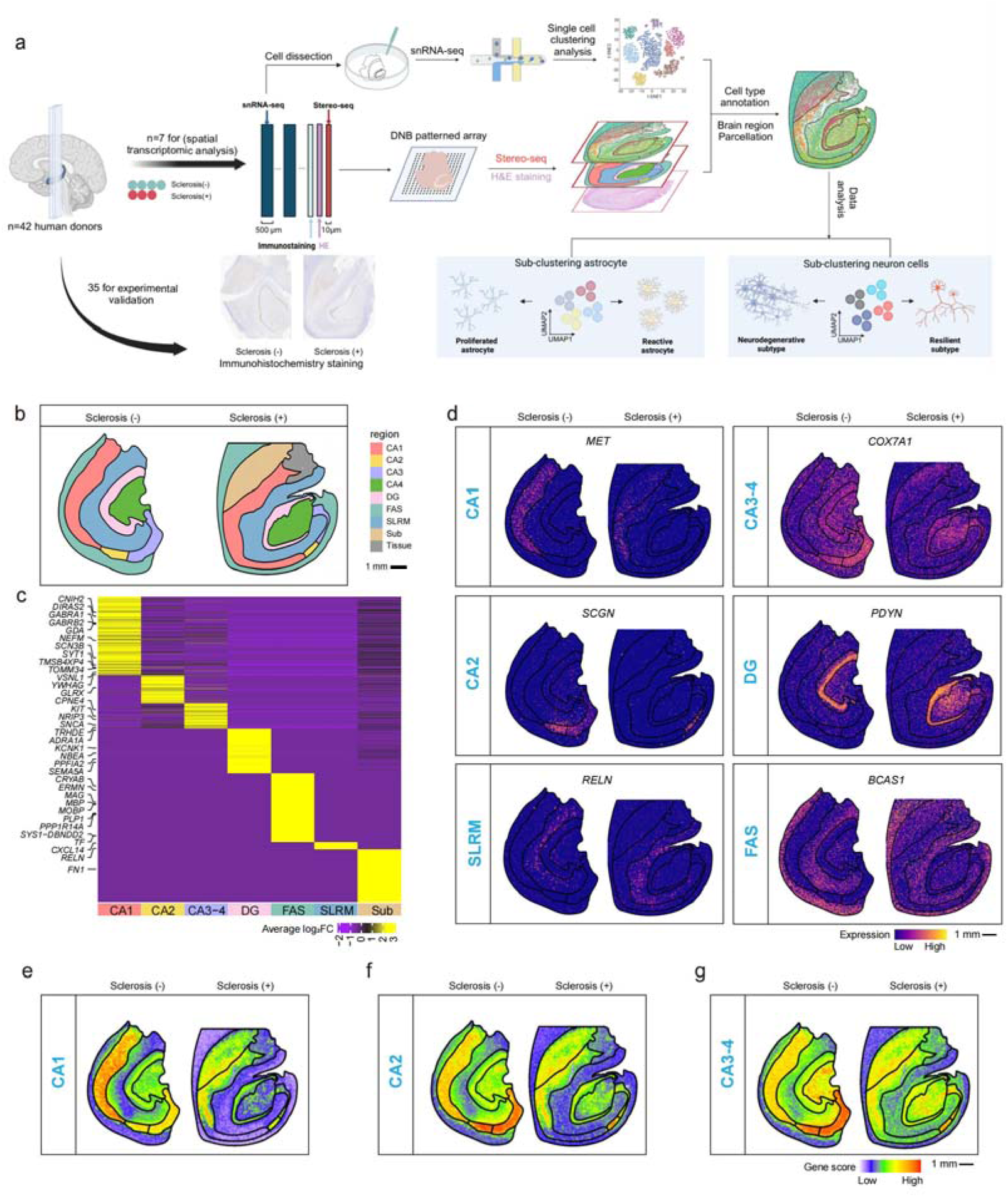
Subregion parcellation of human hippocampus by gene expression pattern. **(a)** An overview of the whole study design. A total of 42 samples with and without HS from intractable epilepsies performed sequencing and staining experiments. Qualified seven samples were performed Stereo-seq and snRNA-seq separately for transcriptomic analysis; and all samples were performed experiments to verify the accuracy of sequencing analysis results. (**b**) Subregion parcellation and annotation of human hippocampus with and without sclerosis. CA: Cornu Ammonis, DG: Dentate gyrus, FAS: fimbria, alveus, and stratum oriens (SO), SLRM (stratum lucidum (SL), stratum radiatum (SR), stratum moleculare (SM)). (**c**) Heatmap shows average log_2_ (fold change (FC)) of subregion-specific expressing genes for each area. (**d**) Spatial expression profile of representative subregion-specific genes. (**e-g**) Spatial gene score profile of DEG modules from (c) between sclerosis (+) and sclerosis (-) in CA1 (**e**), CA2 (**f**), CA3-4 (**g**).

**Fig. 2.**
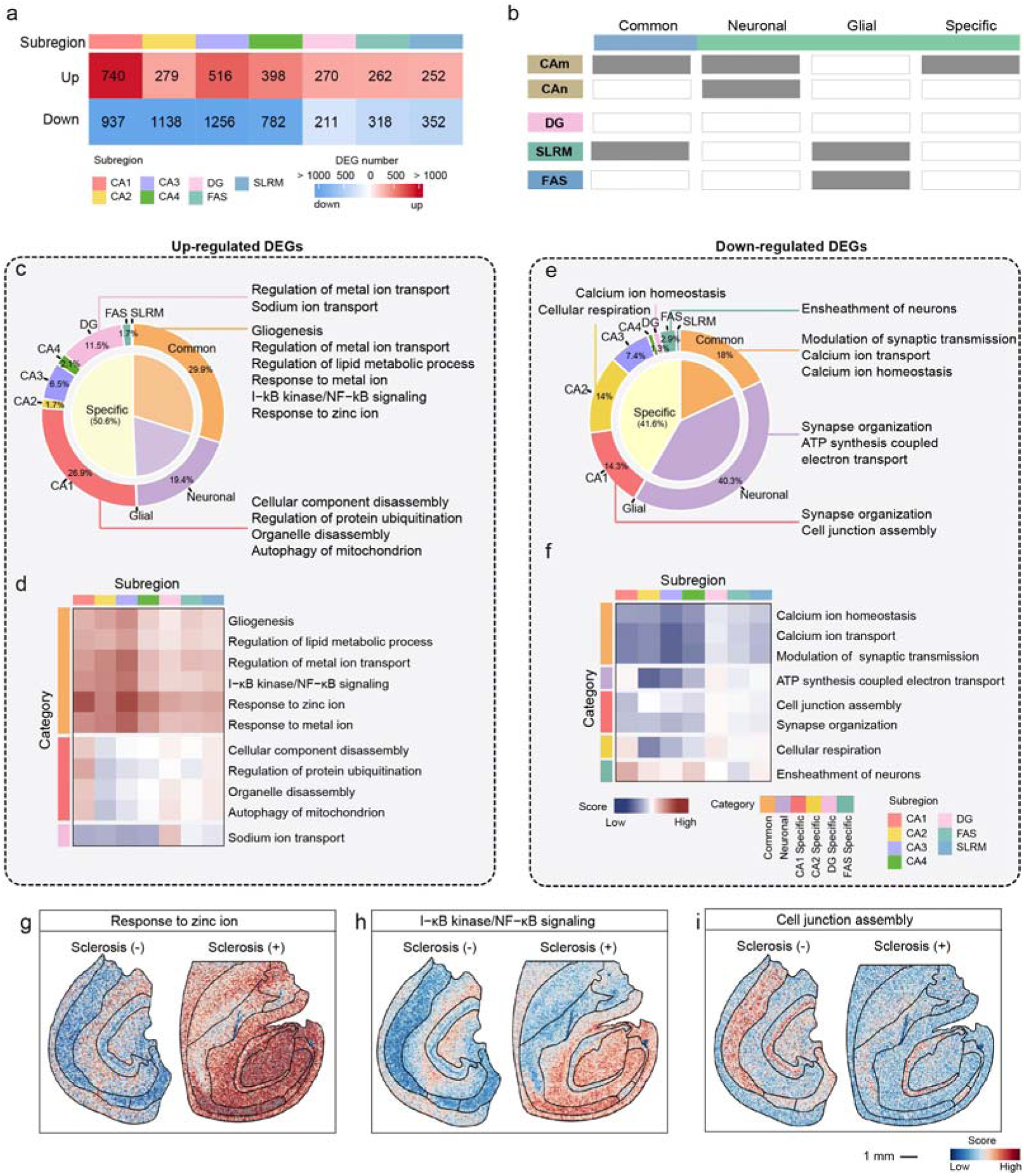
Characteristics and functions of differentially expressed genes in sclerosis in each subregion. (**a**) Heatmap showing the number of DEGs in sclerosis of each subregion. (**b**) Schematic diagram of classification of DEGs in sclerosis of each subregion. CA and DG subregions were categorized as neuronal area, FAS and SLRM subregions were categorized as glial area. Common DEGs (Common): genes in at least one subregion of both neuronal area and glial area; Neuronal DEGs (Neuronal): genes in two and more subregions of neuronal area; Glial DEGs (Glial): genes in FAS and SLRM subregions; Specific DEGs (Specific): genes only in one subregion. (**c and e**) Pie graph showing the proportion of up-regulated (c) and down-regulated (e) DEGs and enriched functions in four categories. (**d and f**) heatmap showing AUCell scores equals sclerosis (+) minus sclerosis (-) of each pathway. (**d**) up-regulated genes enriched pathway. (**f**) down-regulated genes enriched pathways. Red color showed higher scores in sclerosis and blue color showed lower scores in sclerosis. (**g-i**) Scores of pathways in spatial. Response to zinc ion (**g**) and I−κB kinase/NF−κB signaling (**h**) were up-regulated in all brain regions in sclerosis (+). Cell junction assembly (**i**) was down-regulated in the CA1 subregion in sclerosis (+).

### HS-associated changes in gene expression across subregions of hippocampus

To examine the pathological change in sclerosis at transcriptome level, we performed the differentially expressed genes (DEGs) analysis between sclerosis (+) and sclerosis (-) in each subregion. Collectively, we found 7,711 DEGs (2,717 up- and 4,994 down-regulated genes) across seven subregions in HS except for Sub as this subregion was found in limited hippocampal samples. Among the seven subregions with sclerosis, the CA subregions exhibited the most significant changes, especially CA1, with 740 up- and 937 down-regulated genes, and CA3, with 516 up- and 1,256 down-regulated genes (**Fig. 2a, Supplementary Table 7**). Our result implied that CA1 and CA3 were the most vulnerable subregions to sclerosis, but not CA4, which together with CA1 were regarded as pathological landmark of sclerosis^16^. In our analysis, there were 398 up- and 782 down-regulated genes in CA4 with sclerosis, less than that in CA3.

To figure out whether pathological and functional changes in HS were across hippocampal subregions or subregion specific, we first divided up- or down-regulated DEGs into four major categories according to the origin of these DEGs obtained. In hippocampus, CA and DG were considered as neuronal regions whereas FAS and SLRM were recognized as glial regions. DEGs obtained in at least one of neuronal regions and at least one of glial region were categorized as common DEGs. DEGs obtained in two or more subregions of neuronal regions or glial regions, were categorized as neuronal or glial DEGs, respectively. DEGs arised from one specific subregion were categorized as specific DEGs (**Fig. 2b**). Refer to the up-regulated genes in HS, common DEGs (336, 29.9%) were enriched in various functions, such as gliogenesis, lipid metabolic process, metal ion (Zn^2+^) response and transport, and immune response (**Fig. 2c, d, j and h**). As we known, maintaining Zn^2+^ homeostasis was necessary for maintaining the balance between neuronal excitation and inhibition, elevated Zn^2+^ response in HS caused Zn^2+^-induced neurotoxicity contributed to neuronal damage and death associated with seizures^17–19^. The NF-κB signaling pathway exhibited a significant up-regulation trend in all brain regions of the sclerotic samples (**Fig. 2h**). Besides, the abnormal activation of NF-κB signal was closely related to neuronal injury and may be one of the important mechanisms leading to neuronal dysfunction and death. Specific DEGs occupied half (566, 50.8%) of the up-regulated genes, and more than half specific DEGs were CA1-DEGs (302, 26.9%) (**Fig. 2c)**, which were enriched in functions related to cell disaggregation, such as cellular component disassembly, protein ubiquitination, organelle disassembly, mitochondria autophagy (**Fig. 2c and d**). DG-DEGs (129, 11.5%) were enriched in sodium ion transport functions, implying irregular neuronal activity in HS^20^ (**Fig. 2c and d)**. On the other hand, among the 4,994 down-regulated genes, common DEGs (394, 18%) were enriched in functions of synaptic transmission and calcium ion transport (**Fig. 2e and f**). Nevertheless, neuronal DEGs (883, 40.3%) were enriched in synapse organization and ATP metabolism (pronounced in CA2-4) (**Fig. 2e and f**). CA-DEGs, accounting for the largest proportion of specific DEGs, were enriched in functions of synapse organization, cell communication, ATP metabolism. As the vulnerable subregions to HS, neurons in CA subregions were impaired and damaged through the reduction in neuronal connections and the decrease in synaptic density^21^. Down-regulation of cellular connection assembly pathways in CA subregions was contributed to the synaptic deprivation^22,23^. FAS-DEGs were mainly enriched in ensheathment of neurons (**Fig. 2e and f**). We also used public RNA sequencing data of human temporal lobe with MTLE-HS^8^. Up-regulated genes in MTLE-HS were enriched in immune response and gliogenesis, whereas down-regulated genes in sclerosis were enriched in Ca^2+^ and K^+^ transport, cell communication and synapse organization (**Extended data Fig. 3f)**, which was approximately consistent with our results.

In all, based on ranking the number of DEGs, CA was assumed as the most vulnerable region to sclerosis, especially CA1 and CA3. In sclerosis, up-regulated genes were enriched in gliogenesis, metal ion transport, immune response, and lipid metabolic process. Down-regulated genes were enriched in synapse organization, synaptic transmission, cell communication, and ATP metabolism. We speculated these changes of function and gene expression in sclerosis were caused by the cell type composition change and the cell type activities of subregions.

### A single-cell spatial-transcriptional map of human hippocampus with sclerosis

To illustrate the spatial distribution change of cell type composition in sclerosis, we first integrated qualified snRNA-seq data of all samples by using harmony algorithm to remove batch effect^24^ (**Supplementary Table 4**), and annotated the cell types by using canonical cell type markers^14,25^ (**Extended data Fig. 3a-c**). Then we used our previously reported AI-assisted automatic segmentation algorithm (StereoCell)^26^ to delineate cell boundaries based on ssDNA staining (**Extended data Fig. 4a**). On average, we obtained 50,113 segmented cells/sections with 590 genes/cell and 1,227 UMIs/cell after filtering low quality cells (**Extended data Fig. 4b and c**). To annotate the segmented cells in Stereo-seq data, we applied PyTorch-based annotation algorithm (Spatial-ID)^27^ by integrating snRNA-seq with the Stereo-seq data, and the annotated single-cell spatial-transcriptional map (Stereo-map) of human hippocampus with or without sclerosis were constructed (**Fig. 3a**). To validate the cell type annotation accuracy of Stereo-map, we quantified the expression of cell type marker genes in each cell type (**Extended data Fig. 4f**), and then calculated the enrichment of cells surrounding each cell type (**Extended data Fig. 4d**), and computed Pearson correlations to ascertain the correspondence of cell types between the snRNA-seq and Stereo-seq datasets (**Extended data Fig. 4e, Methods**). All these results confirmed the accuracy of cell type annotation on Stereo-map.

**Fig. 3.**
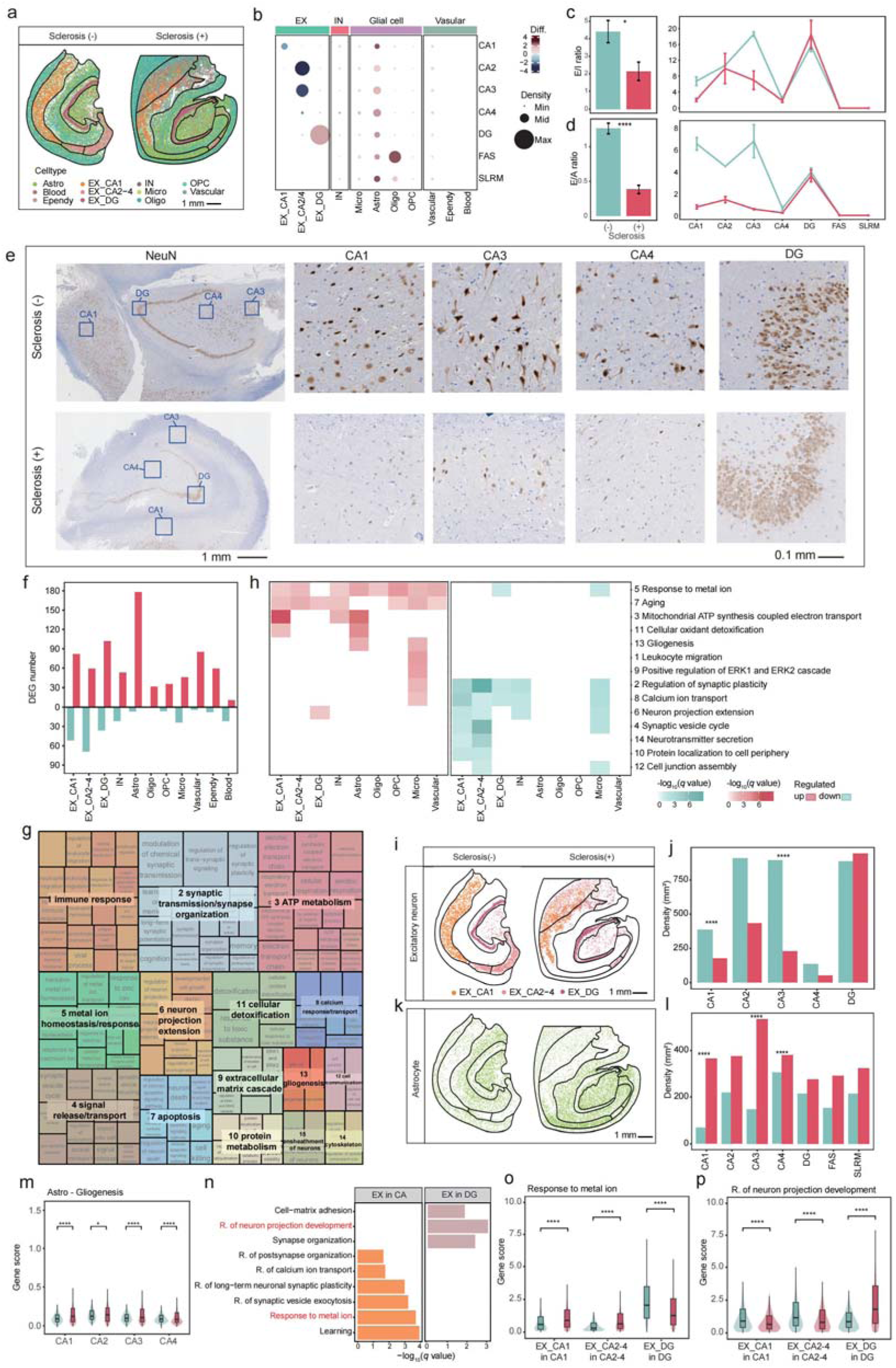
Comparison of single-cell spatial-transcriptomic map of human hippocampus with and without sclerosis. (**a**) Spatial distribution of cell types in hippocampus with and without sclerosis. (**b**) Density change of each cell type in each subregion. The spot size represented average density of each cell type in each subregion over with and without sclerosis groups. The spot color represented an increase (red) or decrease (blue) of density in sclerosis. (**c-d**) Average E/I ratio (**c**) and E/A ratio (**d**) in hippocampus (left) and separated subregions (right) of both with and without sclerosis. E/I ratio: the density ratio of excitatory neurons/inhibitory neurons. E/A ratio: the density ration of excitatory neurons/astrocyte (*t*-test, *\*p* = 0.04*, ****p* < 2e-4). (**e**) NeuN staining of neurons in hippocampus with and without sclerosis. Classical pathologies of CA1, CA4, DG was zoomed in respectively. (**f**) Number of DEGs for each cell type between sclerosis (+) and sclerosis (-). (**g**) Biological pathways overrepresented within cell type-specific DEGs determined by gene set enrichment and semantic clustering analysis of GO terms. Numbers indicated significance rank. (**h**) Aggregate enrichment of GO terms within each biological theme from (i) across cell types by down-regulated (green) and up-regulated (red) DEGs. (**i-k**) Spatial distribution (**i**) and densities of each subregions (**j**) of excitatory neurons in sclerosis (+) and sclerosis (-). (**k-l**) Spatial distribution (**k**) and densities of each subregions (**l**) of astrocyte in sclerosis (+) and sclerosis (-). (**m**) Astrocyte gene scores in the CA1 and CA4 of sclerosis (+) and sclerosis (-) groups for selected GO terms. (**n**) GO terms enriched by excitatory neurons in CA and DG subregions. (**o-p**) Excitatory neuron gene scores in the CA1, CA2-4, and DG subregions of sclerosis (+) and sclerosis (-) groups for two selected GO terms.

Collectively, we identified 14 cell types among a total number of 350,795 segmented cells in human hippocampus, including 151,815 segmented cells with sclerosis, and 198,980 segmented cells without sclerosis. Due to low density of all inhibitory neuron subtypes, we merged them together for analysis at the beginning (**Fig. 3a**, **Extended data Fig. 5**). Inhibitory neurons and most non-neurons were located dispersedly throughout the entire hippocampus whereas various types of excitatory neurons and oligodendrocyte were localized at distinct subregions (**Fig. 3b**, **Extended data Fig. 5**). Cell density change in sclerosis were calculated by comparing each cell type in each subregion between sclerosis (+) and sclerosis (-). As expected, the density of excitatory CA neurons (EX_CAs) were decreased in their located subregions, and the density of astrocyte was increased in all subregions of hippocampus, implying the proliferation of astrocyte in sclerosis (**Fig. 3b**). The result of NeuN staining validated our transcriptomic observations that density of CA neurons was obviously reduced (**Fig. 3e, Extended data Fig. 1e**). Since the density ratio of excitatory and inhibitory neurons (E/I ratio) reflected the brain activities and normal functions, brains with epilepsy were reported to have the imbalanced E/I ratio^28^. Our result indicated that E/I ratio was significantly reduced in sclerosis, especially in CA1 and CA3 (**Fig. 3c**). We also used the density ratio of excitatory neuron and astrocyte (E/A ratio) as index to evaluate the vulnerability of subregions, and the result indicated that E/A ratio was also significantly decreased in sclerosis, with the most notable reduction in the CA1 and CA3 (**Fig. 3d**).

To examine the functional changes of cell types to sclerosis, we performed DEG analysis for each cell type between sclerosis (-) and sclerosis (+). Interestingly, we found that the number of up-regulated DEGs was more than down-regulated DEGs in sclerosis and astrocyte appeared to be dramatically affected by HS as the number of up-regulated genes was predominantly high in this cell type (**Fig. 3f, Supplementary Table 9**). To figure out the important functional change in sclerosis, we performed gene ontology (GO) analysis for all DEGs and semantically clustered the significantly enriched terms into 16 biological themes to aid interpretation (**Fig. 3g, Supplementary Table 10**). Top biological functional themes were consistent with our bin100 data (**Fig. 2c-f**) and reported results^8^, and snRNA-seq data (**Extended data Fig. 3e, Supplementary Table 11**), such as metal ion transport, ATP metabolism, synaptic transmission and organization (**Fig. 3g**). The gene expression score of selected functions in each cell type helped us understand the impaired functions of cells in sclerosis. For example, the expression of genes in calcium ion transport and synaptic plasticity terms were mainly down-regulated in neurons, but that of genes in aging were up-regulated genes in neurons (**Fig. 3h, Supplementary Table 12**). However, the expression of genes in ATP generation was up-regulated in EX_CA1 and astrocyte, which was supposed to be the compensatory mechanism of impaired energy generation. As expected, astrocyte was responsible for up-regulated genes in gliogenesis in most subregions (**Fig. 3h, k-m**), and microglia was responsible for immune response pathways (**Fig. 3h)**.

Although the boundary of DG seemed ambiguous in sclerosis^29^ (**Fig. 3e**), we found that the density of EX_DG was slightly increased in sclerosis (**Fig. 3b, i and j, Extended data Fig. 5**). Since neurogenesis faded away completely by early adolescence in hippocampus, we supposed and found that the hippocampal atrophy caused by MTLE-HS was responsible for the increased density of EX_DG in sclerosis (**Extended data Fig. 4g**), suggesting the resistant character of EX_DG to HS. To compare the character of vulnerable EX_CAs with resistant EX_DG, we obtained DEGs between EX_CAs and EX_DG in sclerosis (-) group (see **Methods**), and GO analysis of these genes indicated that genes associated with the vesicle and ion transport were specifically revealed in EX_CAs neurons, while that associated with cell matrix were specifically found in EX_DG neurons (**Fig. 3n**). Besides, the expression change of genes in particular pathways in sclerosis, such as response to metal ion and regulation of neuron projection development, was opposite in EX_CAs and EX_DG, as the former was up-regulated in EX_CAs and the latter was up-regulated in EX_DG in sclerosis (**Fig. 3o and p**), implying the possible resilient mechanism of EX_DG in sclerosis.

Thus, reduced density of neurons and increased density of astrocytes in CA were the main signature of HS and illustrated the vulnerability of CA subregions in sclerosis, especially CA1 and CA3. Neurons were responsible for dysfunctions of calcium ion transport, synaptic transmission, and activated functions of neuron death and aging, and astrocyte was responsible for up-regulated gliogenesis and apoptosis pathways in sclerosis. The decreased metal ion response and increased neuron projection extension of EX_DG might be responsible for its resistance to HS.

### Spatial distribution change of astrocyte subtypes in sclerosis

To better understand the pathological change in cell types during sclerosis, we further re-clustered each segmented cell type respectively into 2-8 subtypes with distinct markers and biological functions (**Extended data Fig. 6 and 7a, Supplementary Table 13**). Glial and vascular cells were re-clustered into 19 subtypes, and the density of each subtype was changed with various extent across hippocampal regions in sclerosis, especially astrocyte subtypes (**Fig. 4a**). The density of Astro.0 and Astro.1 were significantly increased throughout the hippocampus (**Fig. 4g and j**). To confirm the function of astrocyte subtypes, we searched clues from their marker genes and biological functions (**Extended data Fig. 7a, Fig. 4b**). We found that Astro.0 was highly expressed *SOXD* family genes (**Fig. 4d and h)**, such as *SOX5*, *SOX6*, and *SOX13* (**Extended data Fig. 8a-c**), and genes in Wnt signaling pathway (**Fig. 4c and i**), implying a high association with astrocyte proliferation. Thus, we designated Astro.0 as P_astrocyte (**Extended data Fig. 8m**). The finding of P_astrocyte provided the evidence of astrogenesis in sclerosis^6,7^. Nevertheless, Astro.1, highly expressing genes in inflammatory response like NF-κB signaling pathway and response to oxidative stress (**Fig. 4b, e and k)** was supposed to be reactive subtype, we called it R_astrocyte (**Extended data Fig. 8m**). Genes, highly expressed in disease-associated astrocyte (DAA) from previous reports^30,31^ (**Extended data Fig. 8d**), were also distinctly expressed in R_astrocyte, validating the identity of R_astrocyte (**Extended data Fig. 8m**). In addition, Astro.3, revealed distinct distribution and a clear increase at FAS subregion in sclerosis (**Fig. 4a and l**), was highly expressed genes in pathways of apoptosis such as p38 MAPK cascade (**Fig. 4f and m**), which was possible to be the apoptotic astrocyte (A_astrocyte) whose activation caused the induction of autophagy^32^.

**Fig. 4.**
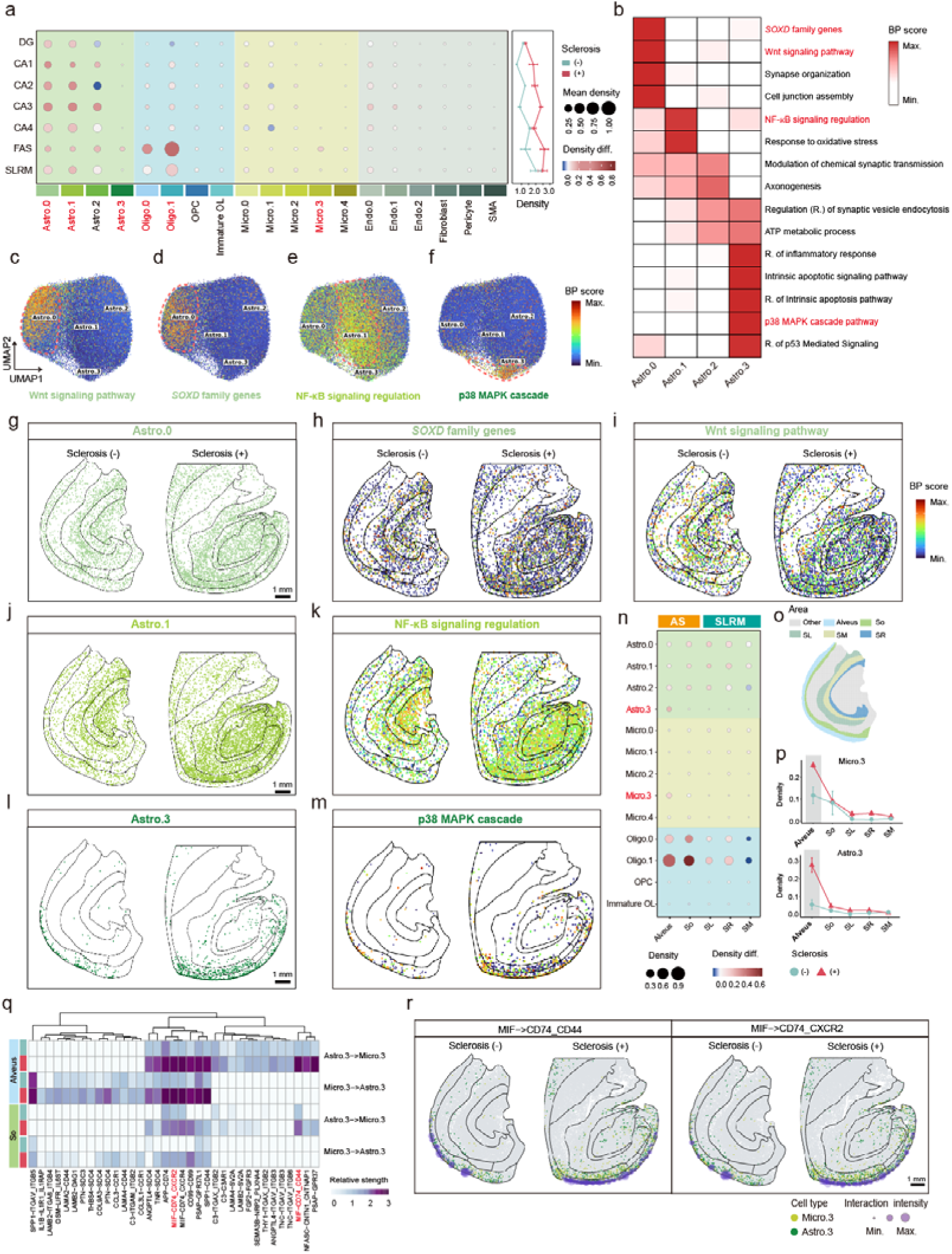
Distribution change of astrocyte across subregions of hippocampus with sclerosis. (**a**) Dotplot showing the density change of each subtype in each subregion. The spot size represented the average density, the spot color represented increase (red) and decrease (blue) of cell proportion in sclerosis, respectively. Density diff., Density difference. (**b**) Heatmap showing expression change of selected enrichment pathways in astrocyte subtypes. (**c-f**) UMAP visualization of Wnt signaling pathway (**c**), *SOXD* family genes (**d**), NF-κB signaling regulation (**e**) and p38 MAPK cascade (**f**) related genes expression in astrocyte subtypes. (**g-m**) The spatial distribution of the corresponding cells and they spatial expression of the pathways highlighted in Figure 4b in hippocampal with (right) and without (left) sclerosis. Scale bar: 1 mm. (**n**) Dotplot showing the density of each subtype change in five small areas of FAS and SLRM. (**o**) The map for small areas parcellation and annotation of FAS and SLRM. SO: stratum oriens, SL: stratum lucidum, SR: stratum radiatum, SM: stratum moleculare. Scale bar: 1 mm. (**p**) Line chart showing cell density of Astro.3 (up) and Micro.3 (down) in each small area with and without sclerosis. (**q**) Heatmap showing the interaction strength of ligand-receptor pairs between Astro.3 and Micro.3 in aleus (blue) and SO (green) with and without sclerosis. (**r**) The scatter plot illustrates the spatially interaction intensity between Micro.3 and Astro.3, as marked in Fig. 3q. The size of the purple circles representing the intensity of the interactions. Scale bar: 1 mm.

Since it was reported that there was a validated interaction between A_astrocyte and reactive microglia (R_microglia) through p38 MAPK casade^33–35^, we wondered whether R_microglia subtype could be found in our study. With our expectation, we found Micro.3, highly expressed genes in inflammatory response (**Extended data Fig. 8e-h and m**), was supposed to be R_microglia whose density was increased distinctly in FAS just like A_astrocyte. Refers to A_astrocyte and R_microglia increased their densities in FAS (**Fig. 4a**), we further subdivided the FAS and SLRM regions into five smaller areas (alveus, SO, SL, SM, SR) based on anatomy and gene expression characters (**Fig. 4o, Extended data Fig. 8j**)^12,15^. The density change of each subtype in each small area showed that A_astrocyte and R_microglia were both distinctly increased at alveus in sclerosis, suggesting the possible interaction between them in alveus (**Fig. 4n-p, Extended data Fig. 8i**). To prove this, we did cell-cell interaction analysis between above two subtypes within 50 µm distance, and found that interaction of ligand and receptor (L-R) pairs with immune functions were increased at alveus in sclerosis, such as MIF-CD74_CD44 (**Fig. 4q and r**), implying further exacerbated the sclerosis-associated pathology within alveus in sclerosis. In addition, the oligodendrocytes also underwent prominent changes in sclerosis. Oligodendrocyte subtypes, Oligo.0 and Oligo.1, functions of myelination^36^ (**Extended data Fig. 8k and l)**, were also showed specific distribution and increased their densities at FAS subregion in sclerosis (**Fig. 4a and n, Extended data Fig. 8l)**.

Overall, the densities of both P_astrocyte and R_astrocyte were increased in sclerosis, especially the area surrounding proximal CA1 subregion. Both A_astrocyte and R_microglia were increased their densities and interactions at alveus in sclerosis.

### Proximal and superficial areas of CA1 were vulnerable to sclerosis

Superficial and deep areas of CA1 were reported to have different sensitivities to sclerosis in epileptic mouse model^9^. To figure out if human hippocampal CA1 had the similar character as mouse model, we used previously reported CA1.Superficial (CA1.sp) and CA1.Deep (CA1.dp) marker genes^37^ for enrichment scoring and effectively distinguished CA1.sp and CA1.dp in human hippocampus (**Extended data Fig. 10a and b, Supplementary Table 14**). In addition, during delineating the CA1 subregions, we noticed obvious difference in CA1 marker gene expression (*MET*) and cell distribution between proximal and distal CA1 of sclerosis (**Fig. 1d, Extended data Fig. 7a, 10c**). To compare the susceptibility of proximal CA1 (CA1.pr) and distal CA1 (CA1.dt) to sclerosis, we parcellated CA1 into CA1.pr and CA1.dt based on the position of vanishing EX_CA1 sharply in sclerosis (see **Methods**) (**Extended data Fig. 10c**). In sclerosis, CA1.pr was highly expressed astrocyte marker genes, such as *GFAP, SPARC, ID3* and *FABP5* (**Extended data Fig. 10d, Supplementary Table 15**), and CA1.dt was highly expressed excitatory neuronal genes, such as *STMN2, NRGN, SNAP25* (**Extended data Fig. 10d**). Due to the heterogenous gene expression and cell composition in sclerosis CA1, we parcellated CA1 subregion into four small areas, CA1_deep_proximal area (CA1.dp.pr), CA1_superficial_proximal area (CA1.sp.pr), CA1_deep_distal area (CA1.dp.dt), and CA1_superficial_distal area (CA1.sp.dt) (**Fig. 5a**). We found that the density EX_CA1 and astrocyte was reduced and increased more severely in CA1.sp.pr, CA1.dp.pr, CA1.sp.dt than CA1.dp.dt area in sclerosis, respectively (**Fig. 5b**), and that genes expression in gliogenesis in proximal CA1 was higher in sclerosis (+) than that in sclerosis (-) (**Extended data Fig. 10f**). Furthermore, the E/A ratio was robustly changed in CA1.dp.pr, CA1.sp.pr, and CA1.sp.dt areas (**Fig. 5c**), illustrating that CA1.sp and CA1.pr were more vulnerable to sclerosis.

**Fig. 5.**
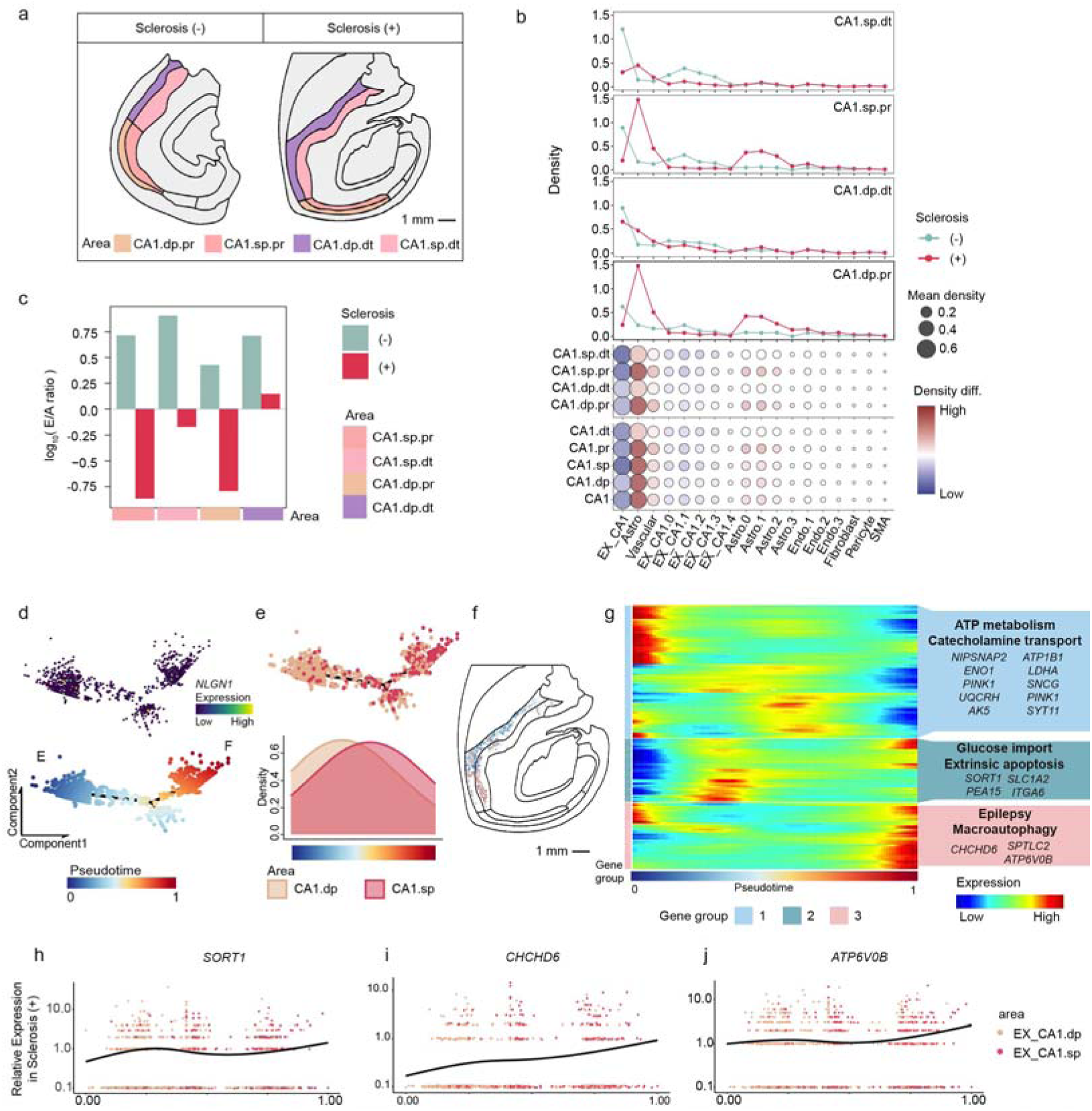
The neuron state change for vulnerable superficial and proximal CA1 of sclerosis. (**a**) The schematic diagram of four small areas for CA1 parcellation. (**b**) Line chart (top) and dotplot (bottom) showing the density and density change of EX_CA1, astrocyte and their subtypes in each area of CA1 subregion. In dotplot, the spot color represented an increase (red) or decrease (blue) of density in sclerosis. Density. diff., Density difference. (**c**) log_10_ (Average E/A ratio) in four small areas of CA1 subregion with and without sclerosis. E/A ratio: the density ration of excitatory neurons/astrocyte. The raw E/A density ratio was adjusted by adding 0.001 to prevent logarithmic singularity due to near-zero denominators. After adjustment, the ratio’s log_10_ value indicates higher EX_CA1 density than Astro if positive, and vice versa. (**d**) Pseudotime analysis of EX_CA1 with sclerosis. Top: pseudotime tree colored by *NLGN* expression. Bottom: pseudotime tree colored by pseudotime. (**e**) UMAP plots were area-colored. Quantitative analysis (bottom) also counted the changes in the percentage of cells in each area over pseudotime. (**f**) Distribution of EX_CA1 colored by pseudotime of (**d**). (**g**) Heatmap showing the expression of pseudotime genes in sclerosis (-) and sclerosis (+) along with trajectory pseudotime time with sclerosis. Each row representing a single gene. Representative GO functions and pseudotime genes of each cluster were list in the right box. (**h-j**) Kinetics plot showing the relative expression of representative genes in sclerosis cluster 3 over pseudotime in EX_CA1 with sclerosis. Points were colored by EX_CA1.sp and EX_CA1.dp.

Since astrocyte and EX_CA1 accounted for the cell composition change in CA1 of sclerosis (**Fig. 5b and Extended data Fig. 10e**), we wondered whether there were changes in the composition and distribution at subtype level. We found that P_astrocyte and R_astrocyte were two of the most dramatically changed subtypes in CA1.dp.pr, CA1.sp.pt, and CA1.sp.dt with sclerosis (**Fig. 5b, 4g and j**). Like the parcellation of CA1.pr and CA1.dt, we also parcellated SLRM and FAS into proximal SLRM (SLRM.pr), distal SLRM (SLRM.dt), proximal FAS (FAS.pr) and distal FAS (FAS.dt), repectively (**Extended data Fig. 10c**).

To dissect the pathological change of EX_CA1, we performed pseudotime analysis^38^ to EX_CA1 subtypes and determined that the beginning of timeline subtype was EX_CA1.3 by using classical synapse marker gene *NLGN1* (**Extended data Fig. 10g-h**), which was running normal neuronal functions, such as synapse organization and synaptic transmission (**Extended data Fig. 7a**). EX_CA1.1 as the terminal subtype with the function of autophagy, was implied as the damaged subtype. Although without distinct spatial distribution pattern for EX_CA1 subtypes, we found that the density of EX_CA1.1 was reduced obviously in CA1.sp.pr, CA1.sp.pr, and CA1.sp.dt areas in sclerosis (**Fig. 5b, Extended data Fig. 10e**). Thus, EX_CA1 neurons passed through normal synaptic transmission, to abnormal energy derivation, and finally into the autophagy process (**Extended data Fig. 10i and 7a**).

Based on the above results that CA1.sp and CA1.pr were vulnerable to sclerosis, and EX_CA1 was vanished a lot in both areas, we speculated that epilepsy-related neuronal autophage might evolve distinctly across areas of EX_CA1, which might be pivotal for cell death program coordinated with hippocampal sclerosis. To extract clues into these transitional states, we used manifold learning leveraged in nearest neighbor information to automatically organize cells in trajectories along a principal tree, reflecting progression of associated biological processes^39^. Reduced dimensional embedding of the automatically learned underlying trajectory produced a spanning tree, showing topological structure of EX_CA1 for sclerosis sample (**Fig. 5d**). *NLGN1* gene was used to confirm the starting site of the pseudo-time, and the result indicated that EX_CA1_sup (EX_CA1.sp) cells reached at the peak in the terminal part of peseudotime in sclerosis sample (**Fig. 5e-f**), implying more severely damaged excitatory neurons in the superficial area of sclerosis. Delving deeper into pseudotime trajectory genes in sclerosis, we found EX_CA1 in sclerosis was impaired at the begining of peseudotime. Genes were mainly enriched in ATP metabolism and catecholimine transport, suggesting the impaired function of energy derivation and increased seizure frequency^40^. Besides the neural dysfunction, we also found neurons passed through glucose import, apoptosis and macroautophage procedure, consistent with the pronounced occurence of EX_CA1 deaths in CA1.sp (**Fig. 5g**). Some of these genes were directly associated with epilepsy, whose expression were increased along with peseudotime in sclerosis, such as *SORT1*, encoded the sortilin transmembrane protein, which acted as a coreceptor with p75NTR for binding proneurotrophins (pro-NGF and pro-BDNF) and inducing apoptosis in neurons^41–43^ (**Fig. 5h**). *CHCHD6*, potential epilepsy-associated gene, increased their expression at the terminal pseudotime to improve cell apoptosis^44–46^(**Fig. 5i**). *ATP6V0B*, encoded a subunit of eukaryotic multisubunit vacuolar ATPase (V-ATPase) enzyme complex, together with *ATP6V0C* were reported as a cause of neurodevelopmental disorder like epilepsy^47–49^ (**Fig. 5j**).

To investigate if there was gradient gene expression and cell type composition changes from proximal to distal of CA1, we subdivided CA1.pr and CA1.dt into 6 continuous groups (**Extended data Fig. 11a**), and screened 804 genes with different expression trend (DEGt) along proximal-distal axis between sclerosis (+) and sclerosis (-) samples by using time-course gene expression method^50^. Then, we clustered DEGt into six clusters with distinct functions (**Extended data Fig. 11b, Supplementary Table 16**). Astrocyte, including all its subtypes, was responsible for C1 cluster, with decreased gliogenesis along proximal-distal axis in sclerosis (**Extended data Fig. 11c, e**), whereas, EX_CA1, including its subtypes, was responsible for C2-4 clusters, with depleted functions of synaptic transmission and energy derivation in CA1.pr of sclerosis (**Extended data Fig. 11c and d**). The density of IN_VIP was higher in CA1.pr of sclerosis, and it was responsible for C5 involved in axon development, implied its neuroprotective function in sclerosis (**Extended data Fig. 11c and f**). In addition to counting the more variable cell types and their subtypes separately, we also counted the E/A ratios of six consecutive groups. We found more significant changes in the proximal three consecutive groups, indicating the greater susceptibility to sclerosis (**Extended data Fig. 11g**). In addition, we also make subdivisions in either FAS or SLRM along the proximal-distal axis with the same methods. As comparing with sclerosis (-), we found that P_astrocyte, R_astrocyte, and R_microglia all exhibited a higher level of cell density in proximal part with sclerosis (**Extended data Fig. 11h-j**).

In all, CA1.sp and CA1.pr were vulnerable to sclerosis with increased gliogenesis and decreased synaptic transmission and energy derivation, which contributed by increased density of astrocyte (mainly P_astrocyte and R_astrocyte) and decreased density of EX_CA1 (including EX_CA1.1).

### Overexpression of extracellular matrix genes was responsible for resilient microenvironment in sclerosis

As CA2-4, especially CA3, were impaired severely in sclerosis (**Fig. 3b-e**), we wondered which subtype of excitatory neuron within CA2-4 changed most in sclerosis. To resolve this, we compared the density of each EX_CA2-4 subtype between sclerosis (-) and sclerosis (+) (**Fig. 6a and b, Extended data Fig. 13a**). Except for EX_CA2-4.3, the density of EX_CA2-4 subtypes was all reduced in sclerosis (**Fig. 6a and b**). Microglia and vascular cells, particularly Endo.1 with function of response to oxygen levels, increased their densities in sclerosis (**Fig. 6a and b, Extended data Fig. 7a**). Like in CA1, the density of P_astroyte and R_astrocyte was also increased in CA2-4, especially in CA3 (**Fig. 6b**).

**Fig. 6.**
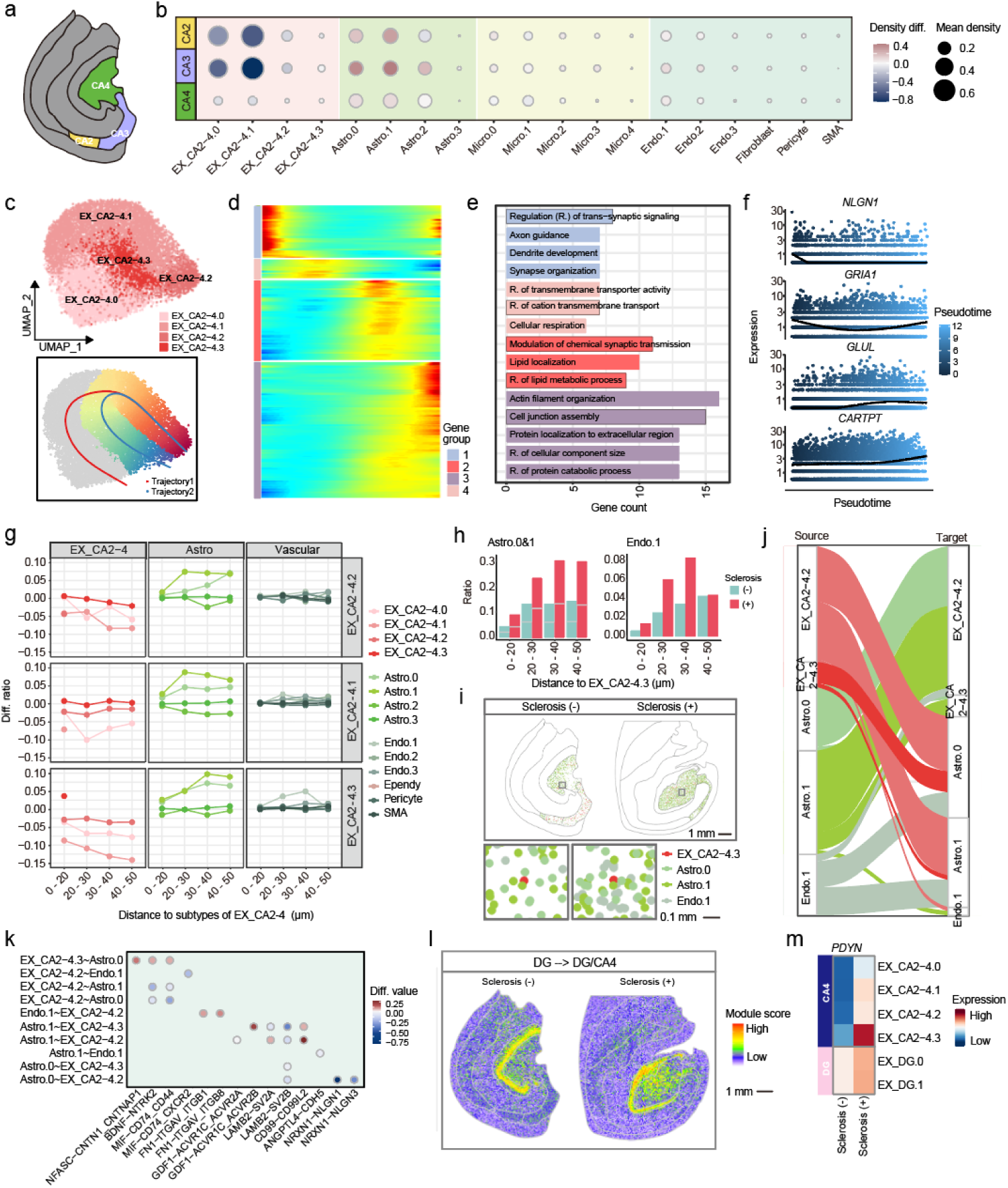
Resilient trajectory of EX_CA2-4 subtypes in sclerosis. (**a**) Hippocampal schematic diagram of CA2-4 subregions. (**b**) The dotplot displaying the density change of each subtype in each subregion of sclerosise the sclerosis (-) group within the subregion. (**c**) Top: UMAP of EX_CA2-4 subtypes. Bottom: UMAP colored by trajectory 2 pseudotime. (**d**) The heatmap showed the expression of pseudotime genes along trajectory 2. The genes were divided into four groups based on their pseudotime expression patterns. (**e**) Bar chart displayed GO functions of gene clusters in (**d**). (**f**) Kinetics plot showing the relative expression of representative genes in cluster 1 and 4 over pseudotime. (**g**) The line graph depicting the proportion difference of each cluster of EX_CA2-4 (left), astrocyte (middle) and vascular cell (right) between sclerosis (+) and sclerosis (-) groups at various distances to each EX_CA2-4 subtype in trajectory 2. (**h**) The bar plot illustrating the proportions of Astro.0&1 (left) and Endo.1 (right) cells within various distance to EX_CA2-4.3 with and without sclerosis. (**i**) The spatial UMAP (top) and enlarged area (bottom) displaying the spatial distribution of Astro.0&1 and Endo.1 surrounding EX_CA2-4.3. (**j**) Sankey diagram illustrating interaction amounts among five subtypes. (**k**) The dotplot showing the differences in the strength of ligand-receptor in subtypes. The colors represent increased (red) and decreased (blue) interaction in sclerosis. (**l**) The spatial UMAP showing the gene module scores of DG-specific genes in the sclerosis (-) group, which were co-expressed in DG and CA4 in sclerosis. Scale bar: 1 mm. (**m**) Heatmap of *PDYN* expression in EX_DG and EX_CA2-4 subtypes.

To investigate if there was a neurodegenerative trajectory in EX_CA2-4 subtypes like EX_CA1, we also performed trajectory analysis^38^ in EX_CA2-4. Using *NLGN1*, EX_CA2-4.2 with the functions of synaptic organization and synaptic transmission (**Extended data Fig. 7a**) was identified as the beginning of the two trajectories: Neurodegenerative trajectory (trajectory 1) (**Extended data Fig. 13b**) and resilient trajectory (trajectory 2) (**Fig. 6c**). Neurodegenerative timeline of EX_CA2-4 was similar, if not totally the same, with pseudotime trajectory of EX_CA1 subtypes (**Extended data Fig. 10g and i**). To illustrate clearly, it was from EX_CA2-4.2 to EX_CA2-4.0 through EX_CA2-4.1 (**Extended data Fig. 13b**). EX_CA2-4.1 and EX_CA2-4.0 were functioned in energy derivation and amyloid-beta formation, respectively (**Extended data Fig. 7a**). The latter was reported to be highly related to cognitive decline caused by epilepsy^51^. To gain deeper insights into the gene expression dynamics along this trajectory, we clustered the numerous trajectory genes, and the result uncovered the gradually functional transition from normal synaptic function, through abnormal energy derivation and aging, to apoptosis and cell death (**Extended data Fig. 13c and d**). For instance, the gene expression levels of *GRIA1* and *RAB3A*, important genes for excitatory synaptic transmission, were reduced along with neurodegenerative timeline. However, apoptotic and aging related genes, *SLC1A2* and *ITM2C*, whose expression level were increased along with neurodegenerative timeline (**Extended data Fig. 13d and e**). Nevertheless, the resilient timeline (line2) seemed to be distinct during sclerosis pathology. It encompassed clusters including EX_CA2-4.2, EX_CA2-4.1 and EX_CA2-4.3 of the pseudotime (**Fig. 6c**). EX_CA2-4.3 was involved in extracellular matrix organization (**Extended data Fig. 7a**), which was similar to the distinct character of EX_DG (**Fig. 3n**). The result of clustering the trajectory genes indicated that genes of group 3 within trajectory were enriched in cell junction assembly, protein localization to extracellular region, and regulation of protein catabolic region (**Fig. 6d and e**), identical to the function of EX_CA2-4.3 (**Extended data Fig. 7a**). *GLUL* and *CARTPT* functioning in strengthen the extracellular matrix^52^ and neuroprotection^53^, were highly expressed at the terminal pseudotime (**Fig. 6f**).

To quantify changes of cell composition around subtypes of EX_CA2-4, we set 4 concentric circles spaced apart with each subtype of EX_CA2-4 at the center. The results showed that the proportions of non-neuronal cell type were increased by distance of each EX_CA2-4 subtype, such as astrocyte and vascular subtypes (**Extended data Fig. 13f**). To compare the microenvironment of resilient subtype with and without sclerosis dynamically, we calculated the difference of proportion between samples with and without sclerosis of each cell type at each distance of microenvironment (see **Methods**). The result revealed that the proportions of P_astrocyte and R_astrocyte were increased abruptly and then kept at high level surrounding in both EX_CA2-4.2 and EX_CA2-4.3 microenvironment (**Extended data Fig. 13g, Fig. 6g-i**), however, the proportion of Endo.1 was particularly increased in EX_CA2-4.3 microenvironment, but not other EX_CA2-4 subtypes (**Extended data Fig. 13g, Fig. 6g-i**). To examine the cell communications surrounding resilient microenvironment, we performed cell-cell interaction analysis within 50 μm microenvironments of EX_CA2-4 subtypes by using StereoSiTE^54^. We observed significant changes of interaction intensities in sclerosis, with enhanced interactions among non-neuron subtypes, especially astrocyte and endothelial cell, and reduced interactions among neurons subtypes (**Extended data Fig. 13h**). Focusing on EX_CA2-4.2 and EX_CA2-4.3 in resilient trajectory, we found that EX_CA2-4.2 had more interactions than EX_CA2-4.3. EX_CA2-4.3 with small number interaction pairs, had more interactions with P_astrocyte, and fewer number of interactions with R_astrocyte, whereas, EX_CA2-4.2 had similar amount of interactions with P_astrocyte and R_astrocyte, and a few interactions with Endo.1 (**Fig. 6j**). The distinct interaction between EX_CA2-4.2 and Endo.1 implied that it was pivotal for neuron activities, which explained that complementary strategy of increased Endo.1’s density in the EX_CA2-4.3 microenvironment because of depleted interaction between them. It was reported that enhancement of BDNF and its interaction to NTRK would reduce 80% frequency of seizure^55^, Interaction strength of BDNF-NTRK was decreased between EX_CA2-4.2 and P_astrocyte/R_astrocyte, however, it was increased between EX_CA2-4.3 and P_astrocyte (**Fig. 6k**), implying the potential resilient clues. Besides, loss of function of NFASC was reported to cause epilepsy^56^. The interaction between NFASC and its receptors CNTN1 was strengthened in sclerosis in EX_CA2-4.3 microenvironment (**Fig. 6k**), implying the resilience of EX_CA2-4.3 in sclerosis.

Since extracellular matrix activities were both enriched in EX_DG and EX_CA2-4.3, we wondered if there was something in common between EX_CA2-4.3 and EX_DG in sclerosis. As previous studies reported that the boundary of DG subregion was ambiguous in HS, which may be contributed by the softened extracellular matrix environment and granule cell dispersion^57,58^. We parcellated DG subregion into three small areas based on classical anatomy and gene expression characters: molecular layer of DG (MLDG), granule layer of DG (GLDG), and pyramidal layer of DG (PLDG) with area-specific expressed genes^12,15^ (**Extended data Fig. 14a**), such as *SCL1A2* for MLDG, *KCNIP4* for GLDG, and *GAD2* for PLDG. We also found that MLDG and PLDG shared common genes such as *AQP4,* indicated similar gene expression profile of the two areas (**Extended data Fig. 14b and c**). The density of EX_DG was increased in MLDG in sclerosis, contributed mainly by EX_DG.0 which was involved in energy derivation (**Extended data Fig. 7a, 14d-f**). GO functional analysis of up-regulated DEGs in sclerosis of EX_DG.0 revealed that highly expressed genes in sclerosis were related to lipid metabolism regulation, autophagy processes, and apoptotic signaling pathways (**Extended data Fig. 14g**), potentially reflecting sclerosis pathology caused by epilepsy. Besides, we found several genes (*ADRA1B*, *NRN1*, *BDNF* and *PDYN*), selectively expressed in DG in normal hippocampus, exhibited an expanded expression to adjacent CA4 subregion in sclerosis (**Fig. 6l, Supplementary Table 17**). Taking *PDYN* as an example, it was highly expressed in EX_CA2-4.3, much higher than EX_DG.0 and EX_DG.1, although increased its expression in later subtypes, in sclerosis (**Fig. 6m**), implying the potential significance of extracellular matrix-related pathways in resistant to pathological process of sclerosis.

In summary, we found a potential resilient mechanism related to extracellular matrix organization from resilient trajectory of EX_CA2-4 subtypes, the strengthened interactions of PDYN-NTRK, NFASC-CNTN1 between astrocyte and resilient subtype might be important for resilience to HS, providing potential clues for surviving cells from sclerosis.

### Spatial distribution change of inhibitory neuron subtypes in sclerosis

The proportion of inhibitory neurons was quite low in human hippocampus (**Extended data Fig. 9**), and four types of inhibitory neurons revealed different spatial distribution patterns within hippocampus (**Extended data Fig. 15a and b**). Simply, the densities of IN_SST and IN_VIP were relatively higher than IN_PVALB and IN_LAMP5 in sclerosis, especially in the CA1 region which was probably caused by hippocampal atrophy in sclerosis (**Extended data Fig. 4g, 15a and b**). IN_SST was primarily located at the boundaries between CA1 and SO, and between CA4 and PLDG, whose density was increased in former and decreased in latter boundaries (**Extended data Fig. 15b**). Besides, the principle subtypes, SST.0 and SST.1, seemed to contribute the density changes of IN_SST in sclerosis (**Extended data Fig. 15a and 16**). Nevertheless, IN_VIP was globally distributed in hippocampus and showed increased density in sclerosis (**Extended data Fig. 15a and b**). The density of VIP.0, the principle subtype of VIP neurons with distinct function of amyloid fibril formation (**Extended data Fig. 7a**), was increased in DG, CA1 and SLRM with sclerosis (**Extended data Fig. 15a and c**) consistent to previous report showed that intractable epilepsy had an increased burden of soluble Aβ pathology through experimental studies^51^ Amyloid beta precursor protein, *APP*, whose expression was increased in VIP.0 of sclerosis, and so did its receptor *CD74* in responding cells (**Extended data Fig. 15d and e**). On another hand, previous studies also reported that *VIP*, as a neuroprotective neuropeptide^59^, exerted potential anti-inflammatory effects^51^, enhanced neurogenesis^60^, and contributed to maintain the integrity of the blood-brain barrier^37,59,61^ in epilepsy. Then we wondered which VIP subtype was highly expressed *VIP* and its receptors. We found that *VIP* and *VIPR2* were both highly expressed in VIP.5 (**Extended data Fig. 15f and g**) functioning in catecholamine secretion (**Extended data Fig. 7a**), which was consistent with the previous report that VIP peptide helped to stimulate catecholamine secretion^29^ whose elevated expression provoked epileptic activity^40^. Besides, our results also revealed that the expression *VIP* receptors, *VIPR1* and *VIPR2*, but not *ADCYAP1R1*, were elevated in EX_DG (especially DG.1) and immature oligodendrocyte in sclerosis, respectively (**Extended data Fig. 15c-e**). In total, IN_VIP possibly acted bidirectional functions in sclerosis.

## Discussion

We reported here a single-cell level and genome-wide spatial transcriptomes of human hippocampus with and without sclerosis. We subdivided subregions of hippocampus; molecularly defined the cell types and subtypes of human hippocampus; compared their composition differences by subregions in MTLE with and without HS. We revealed that CA1 and CA3 were more vulnerable than other subregions to MTLE-HS, with signature of decrease in excitatory neuron and increase in astrocyte. These pathological characters were more obvious in proximal superficial CA1 subregion. Refer to astrocyte, we observed P_astrocyte and R_astrocyte were increased their densities to CA subregions where excitatory neurons disappeared. A_astrocyte increased its interaction with R_microglia in alveus of sclerosis, implying the robust inflammatory activity of alveus were in sclerosis. Excitatory neurons undergone synaptic impairment, energy dysfunction, aging until death process in sclerosis. Importantly, we found a resilient subtype, EX_CA2-4.3, highly expressed extracellular matrix related genes including *PDYN*, and was increasing the interaction of BDNF-NTRK and NFASC-CNTN1 to withstand the damage from sclerosis.

To our knowledge, this is the first report to systematically elucidate the subregional pathological change of gene expression and cell type composition in human MTLE-HS. In this study, our study provides new insight on how anatomical structure, or the spatial location of cells, influences cells’ vulnerability to disease, indicating that anatomical structure determines not only organ functions, but also vulnerability of cells where located in. Taken CA1 subregion as an example, excitatory neurons in sclerosis CA1, went through synaptic impairment, energy dysfunction, aging to death procedure, however, both CA1_proximal and CA1 superficial area seem complete the neurodegenerative procedure much earlier than CA1_deep area. The susceptibility of CA1 and CA3 to sclerosis might be related to reliability of the blood supply, given the reason that only a long-branched blood vessel of anterior choroidal artery supplies the blood to CA1 and CA3, which has lower hypoxia resistance than mixed blood vessels^62^.

By using mouse model of hippocampus sclerosis, impaired CA1 neurons in CA1_superfical area displayed neurodegenerative transcriptional trajectories, apoptotic related genes were highly expressed in the epileptic neurons^9^. Our study provides more evidence that impair excitatory neurons in CA subregion, both EX_CA1.1 and EX_CA2-4.0 were enriched in expressing neuron death and apoptotic genes. Such as *CHCHD6*, encoding a core protein of the mammalian mitochondrial contact site and cristae organizing system^63^, increased its expression to improve cell apoptosis in epileptic pathological process^45,46^, and was also an aging factor^64^ and directly connected Aβ processing in neurodegenerative disease-Alzheimer’s disease (AD)^63^, suggesting the common clue between epileptic processing and aging. Protein phosphatase 3 catalytic subunit A (*PPP3CA*), encodes catalytic subunit of calcium-calmodulin-related serine-threonine phosphatase, related to epilepsy^65^ and age-related macular degeneration^66^, implies the similarity of impaired excitatory neurons in epilepsy and aging. As well, a report indicated that a list of psychiatric disease related genes, share with aging, mainly reflect nurture factors such as socioeconomic and behavioral changes associated with living with the disease, environmental exposure and medication^67^. In addition, neurodegenerative diseases, such as Alzheimer’s disease (AD) and Parkinson disease, were highly correlated with aging process^68–70^. However, a recent study report that impaired neurons in the cerebral cortex of schizophrenia (SCZ) display neurodegenerative or aging properties^71^. SnRNA-seq of human prefrontal cortex from 191 human donors indicated that a group of synaptic transmission related genes decline with age and SCZ, complement component 4 (C4), classical aging marker, highly expressed in astrocyte of SCZ donors’ brain. Perineuronal net abnormality is also shared in aging and neurological disease, such as autism, schizophrenia, as well as Alzheimer’s disease^72^. Based on the above discoveries, we speculated the possibility of that disease or complicated environmental or social factors speeding up some of neurons in the distinct focus go through aging process, so that psychiatric disorders occur at any age. The neural senilism hypothesis is needed to be further discussed systematically.

Beyond the neurodegenerative procedure, some of the neurons go through resilience pathway, such as EX_DG, EX_CA2-4, IN_SST, and IN_VIP. To be pointed that EX_CA2-4.3, a small number group cells with similar characters of EX_DG, reveals no obvious density change in CA2-4 subregions of sclerosis, implying the resilient ability to sclerosis. EX_CA2-4.3 highly expresses genes in protein localization to extracellular matrix and genes in axonogenesis, suggesting the dose compensation to the softened extracellular matrix caused by sclerosis. EX_CA2-4.3 also expresses a list of DG marker genes of hippocampus without sclerosis. *PDYN*, encodes prodynorphin which is a precursor to dynorphin opioid peptides, is only expressed at DG subregion of hippocampus, but we observed it highly expressed in EX_CA2-4.3, suggests the epigenetic level has been changed in MTLE-HS. Reports show *PDYN* is implicated in brain/mental disorders^73^, such as drug addiction, Alzheimer’s disease and epilepsy^74^. Knock out *PDYN* expression in cortical cortex of mouse displays deficiency of extracellular signal-regulated kinase (ERK1/2)^75^, implying the pivotal function in extracellular signaling of *PDYN*.

The neuropeptide level changed in the epilepsy brain^76^. We first observe the density of IN_VIP and IN_SST increases in CA1 in HS because of hippocampal atrophy in HS^77^.VIP, known as a neuroprotective peptide^78^, is reported to have suppressive effects on absence seizures through inducing transformation of glutamate to GABA and increasing GABA concentration^79^. Besides, the increased interaction between *VIP* and its receptors in seizure focus implied a mechanism of survivability when facing the increased excitation for injury in a seizure focus^80^. However, the function for *VIP* inducing catecholamine secretion was validated *in vivo* years ago^29,81,82^, highly secreted catecholamine aggravates seizure susceptibility^40,83^, suggesting the side effect of increased VIP in epilepsy, our study indicated both protective and susceptive side of VIP in sclerosis. Taking together with neuropeptide SST, *in vivo* studies in rodents reveals that *SST* and its receptors mediates the antiepileptic actions in hippocampus^84–86^, *in vitro* studies shows that SST dysfunction accounts for epileptogenesis and susceptibility, suggests the antiepileptic function of SST although lacking human tissue experiments^87^. Our data provides the transcriptional increase of *SST* expression and antiepleptic character of IN_SST in the distal and deep area of CA1 in sclerosis, however, the role of neuropeptide SST and VIP and if they can be therapeutic targets need to be further validated.

Energy derivation is seriously impaired in sclerosis caused by epilepsy^88^. As we known, astrocyte serves as the intermediation between vascular and brain neurons^89^. It combines with vascular cells, like endothelial cell and pericyte, and neurons to form neurovascular unit (NVU) by tight contact of endfeet and synapse^90^. Nutrients like glucose and oxygen get through NVU and into the neurons, and metabolites like lactic acid and toxics go through NVU into the vessel^91^. Thus, the well functioned neuron and astrocyte system is quite important for the brain function^92^. Dysfunction of energy derivation reveals the impairment of neuron and astrocyte system running^92^. Reports have indicated that NVU integrity and synaptic plasticity drive alternations of energy metabolism in human brain, and abnormal NVU activation is observed in sclerosis^93,94^. Our study observed increased density of reactive astrocyte and endothelial cell, implied the abnormality of NVU in sclerosis. However, the increase of NVU in epilepsy playing consequential role or complementary role needs to be further investigated.

Although with original and oriental findings, there were still several limitations in our study. For example, the inherent individual differences among donor samples, such as different types of sclerosis, epileptic causes, seizure history, pathological stages of epilepsies, medications are all possible factors to affect gene expression profile and cell type composition change in sclerosis. Besides, the sample size needs to be increased, though we collected for years from surgery donors to verify our results. In addition, our samples were mainly from donors of Han nationality of China, which may reflect the characteristic and mechanistic crew of hippocampal sclerosis caused by epilepsy, however, if it is universal to human being needs to be verified by other nations, such as Caucasian and African.

In all, we first drew single-cell resolution spatial transcriptomic map of human hippocampus with and without sclerosis. CA1 and CA3 were two severely impaired subregions in sclerosis. Refer to CA1, proximal and superficial areas were more vulnerable to sclerosis than CA1.dp areas. P_astrocytes and R_astrocytes were filled in where excitatory neurons disappeared. Excitatory neurons process two state fates, neurodegenerative fate and resilient fate, the latter provides potential clinical clues for epilepsy therapy. The whole study offering a new insight into understanding the pathological mechanism of sclerosis.

## Methods

### Ethics statement

All patients and their guardians who participated in this study signed a written informed consent form, including one that permitted data sharing. All experiments involving human brain tissue in this study were approved by the Human Ethics Review Committee of West China Hospital, Sichuan University (Research Ethics Approval Number: 2022-1159) and the Institutional Review Board on Ethics Committee of Beijing Genomics Institute (BGI) (Research Ethics Approval Number: BGI-IRB 22179).

### Prospective inclusion of clinical patients

Prospective inclusion was made of patients scheduled for surgical resection of refractory temporal lobe epilepsy foci at the Department of Neurosurgery, West China Hospital, Sichuan University, from December 1, 2021, to October 10, 2024. The inclusion criteria for this study were as follows: 1) The clinical diagnosis of epileptic seizures and syndromes refer to the standards of the International League Against Epilepsy (ILAE)^95–97^. 2) Age between 5 and 45 years, with no gender restrictions; 3) A clear clinical history, epileptic symptoms, at least 72 hours of video electroencephalogram (EEG) during the ictal and interictal periods, and cranial MRI examination results; 4) Temporal lobe epilepsy symptoms with EEG showing unilateral temporal lobe onset.

The diagnosis of non-lesional TLE was based on the following criteria: 1) MRI showed no other organic lesions except for hippocampal atrophy; 2) Focal ictal TLE patterns recorded by scalp and depth electrode EEG; 3) EEG records at least three spontaneous seizures, with no independent contralateral epileptic seizures; 4) Clinical features consistent with TLE seizures. When the EEG and MRI examinations showed inconsistent lesion origins or MRI showed multiple lesions, a PET scan must be performed.

Once the patients completed the above auxiliary examinations, a comprehensive team consisting of neurologists, neurosurgeons, radiologists, and EEG room doctors from West China Hospital, Sichuan University, conducted a multidisciplinary consultation. The preoperative assessment included the patient’s clinical history, physical examination results, epileptic seizure symptoms and severity, imaging data, and electrophysiological data. The surgical plan was tailored to the patient’s individual needs and epileptic symptoms in a multidisciplinary team meeting. When all preoperative information pointed to a consistent unilateral focus of focal epileptic seizures, the patient could directly undergo epilepsy-focus resection surgery after multidisciplinary expert consultation. If positive findings were present in the temporal lobe of the patient’s preoperative MRI, the resection range was determined in combination with the extent of the lesion and electrophysiological data. When there were no clear positive findings in the preoperative MRI, the resection range was determined based on clinical electrophysiological data and invasive EEG monitoring results. The extent of the temporal lobe resection was also considered the severity of epilepsy and the potential risk of complications; generally, the range on the left temporal lobe was 4.5-5 cm from the temporal pole, and on the right temporal lobe, it was 5-5.5 cm from the temporal pole.

The exclusion criteria for this study were as follows: 1) the presence of other neurological diseases such as neuropsychiatric diseases, cerebrovascular diseases, intracranial tumors, or congenital vascular malformations; 2) history of craniocerebral trauma; 3) history of cranial surgery; 4) existence of severe neuropsychiatric disorders; 5) a personal history of any mental or substance use disorders; 6) patient or representative refuses to participate in this study.

### Collection of clinical data

Researchers prospectively collected clinical data through face-to-face interactions for TLE patients meeting the criteria above. The current medical history includes the age of onset of epileptic seizures, the duration of the epilepsy condition, the age at surgery, and the frequency of seizures, with the latter being determined by the average frequency of seizures in the three months before surgery, given the irregularity of seizures in some patients. The past medical history included information on whether there was a history of febrile convulsions, or early diseases of the patient (such as difficult birth, high fever in childhood, encephalitis, trauma, etc.). The form of patient seizures, including whether they involve secondary generalized tonic-clonic convulsions, was also recorded as a separate item.

### Surgical specimen sampling

All surgeries were conducted strictly for therapeutic purposes. We prospectively included surgically resected hippocampal and temporal lobe specimens from patients undergoing temporal lobe epilepsy focus resection surgery. All patients were treated by the same neurosurgeon, who performed the en bloc resection without cauterisation from the posterior margin to the uncus. Following the removal of the epilepsy focus, a temporal lobe specimen and a part of the hippocampal tissue were collected. The hippocampus was sectioned along its longitudinal axis. Half of the hippocampal tissue was placed in 4% paraformaldehyde and fixed in a 4 °C refrigerator for at least 24 hours, after which it was used for routine pathological analysis. The remaining half of the hippocampal tissue was carefully and gently blotted with lint-free paper to remove moisture from the surface of the specimen. It was then embedded in the Optimal Cutting Temperature (OCT) compound according to the sampling order. The embedding box was rapidly frozen in isopentane pre-cooled with liquid nitrogen, and the freezing of the sample was completed within 3 minutes. The entire process was finished within 30 minutes after the specimen was excised. The frozen samples were transferred to a -80 °C freezer for storage until further experimental use.

### Paraffin embedding and sectioning

The surgical brain specimen was carefully removed from the fixative, and the sample surface was rinsed with tap water to remove the fixative. The brain tissue sample was dehydrated through a graded series of alcohol concentrations: 50%, 70%, 85%, 95%, and anhydrous ethanol (100%), gradually removing water from the tissue block. The brain tissue was immersed in a mixture of xylene and paraffin at a volume ratio of 1:1 for 1 hour, and then the sample was processed with paraffin for 2 hours. After paraffin infiltration, the sample was placed in an embedding box, molten paraffin was gently poured over the sample, and the paraffin was allowed to cool and solidify in a -20 °C freezer. The paraffin sectioning machine was adjusted, the cooled paraffin block was removed and trimmed, and the block was attached to the sectioning machine. The advancer and wheel were adjusted to section the brain tissue sample at a thickness of 5 μm along the coronal plane. A few drops of double-distilled water were placed on a microscope slide. The slide was used to pick up the floating paraffin sections in the 40 °C warm water of the section stretcher, ensuring that the sections were fully extended. Filter paper was used to absorb the moisture from the sections. Then, the slides were placed in an oven to dry for 30 minutes, and the sections were stored at room temperature for later use.

### H&E staining of paraffin sections

The paraffin sections were cleared with two changes of xylene, each for 20 minutes, and then hydrated through a series of alcohol concentrations: anhydrous ethanol (100%), and 75% alcohol, each for 5 minutes. The paraffin sections were then soaked in double-distilled water for 3 minutes. The sections were stained with hematoxylin for 3 to 5 minutes, washed with double-distilled water for 15 minutes, and soaked in a 1% hydrochloric acid solution for 10 seconds until they turned red. Once the sections turned red, they were removed and placed in an ammonia solution until they turned blue. The sections were then rinsed under running water. The paraffin sections were dehydrated with 85% and 95% alcohol solutions for 3 to 5 minutes each, and finally soaked in a 0.5% eosin solution for 5 minutes. The stained sections were placed in anhydrous ethanol for 3 to 5 minutes, then in xylene for 2 to 5 minutes before being removed to absorb the liquid on filter paper. Neutral resin was added to the center of the section, and it was carefully covered with a cover slip, avoiding the formation of air bubbles. The H&E stained hippocampal region of the brain tissue was observed under a fluorescence microscope imaging system in bright field conditions. Image-Pro Plus 6.0 software was used to analyze the histopathological changes in the H&E stained tissue.

### Pathological diagnosis and annotation of surgical specimens

Surgical samples were independently diagnosed by two expert pathologists from the Department of Pathology at West China Hospital, Sichuan University, based on H&E stained sections of surgical hippocampal and temporal lobe specimens. If a definitive diagnosis cannot be made solely based on H&E staining, special staining or immunohistochemical staining was performed according to the pathological diagnostic process established by the European Neurology Society to assist in the diagnosis. The diagnosis of hippocampal sclerosis was made according to the 2013 ILAE criteria^16^, which state that the affected hippocampus could be visibly hard and reduced in volume, with corresponding expansion of the temporal horn of the lateral ventricle; 1) Type I HS ILAE classification (classic hippocampal sclerosis) refers to the loss of neurons in the CA1, CA4/end folium, and CA3 regions (sparing CA2), associated gliosis, dispersion of granule cells in the DG, and scattered enlarged neurons in the CA4/hilus; 2) Type II HS ILAE classification (CA1 hippocampal sclerosis) referred to severe neuronal loss and significant gliosis in the CA1 region; 3) Type III HS ILAE classification (End-folium hippocampal sclerosis) referred to significant neuronal loss and gliosis in the CA4 region and/or end folium; 4) Type IV HS ILAE classification: there was no significant loss of neurons in the CA1-4 regions, and the granule cell layer in the DG was normally arranged or (locally) mildly disordered (**Supplementary Table 1**). All diagnoses were made by two pathologists, one of whom did not participate in the initial diagnosis. If there was disagreement between the two, the surgical sample in question was excluded.

### Cryosection and H&E staining

The cryostat was adjusted, the OCT-embedded tissue block was removed, trimmed, and the frozen tissue block was secured to the sample holder. The advancer and wheel were adjusted to section the hippocampal tissue sample at a thickness of 10 μm along the coronal plane. After obtaining complete, wrinkle-free, and undamaged frozen sections, the sections were fixed with 95% ethanol for 6 to 10 seconds. The slides were immersed in a hematoxylin-stained jar for 3 to 5 minutes and then rinsed in water for 3 to 5 seconds. Subsequently, the slides were immersed in 1% hydrochloric acid ethanol for 3 seconds and rinsed in water for 1 to 2 seconds. At this point, under a microscope, it was observed that if the cell nuclei were purple-blue and the cytoplasm was colorless, the operation was correct. The nuclei turned blue in 5 to 10 seconds using the bluing solution (1% ammonia water); they were rinsed briefly in water for 1 to 2 seconds under the microscope, and if the nuclei were blue, the operation was confirmed to be correct. The sections were moistened with 75% ethanol. The slides were immersed in an eosin (alcohol-soluble) staining jar for 30 to 60 seconds and then thoroughly rinsed with water. Under the microscope, it was confirmed that if the nuclei were blue and the cytoplasm was red, the operation was correct. The sections were cleared with 95% alcohol to extend their preservation time. Under the microscope, the cell nuclei appeared blue and the cytoplasm appeared pink. The sections were air-dried, neutral resin was added, and they were covered with a cover slip to complete the staining process.

### RNA integrity quality control and completeness of histological sampling site quality control

Frozen sections of 10 μm thickness were cryosectioned at -18 °C in a Leica CM1950 cryostat and placed on aseptic positively charged anti-stripping glass slide. To ensure high-quality samples, 20 consecutive 10 μm sections were cut, or until the frozen tissue block surface was complete, and placed in low adhesion tubes (Eppendorf, ep) tubes. A kit was used to test the RIN of a portion of the donor sample block (**Supplementary Table 3**). In the stereo-seq experiment, RIN > 6.0; sampling included a complete DG area; the sampling site was free of obvious ice crystals to meet the quality control requirements of this study. The 10 μm sections were affixed to standard microscope slides and stained with H&E for orientation and histological structure integrity quality control. Hippocampal samples with intact dentate gyrus and hippocampal CA regions removed surgically were selected for further experiments.

### Stereo-seq chip preparation

The capture chips were fabricated following the Stereo-seq protocol^11^. Initially, to establish the DNB array for in situ RNA capture, we synthesized oligonucleotides featuring a randomized 25-nucleotide coordinate identity (CID). These oligonucleotides were then circularized using T4 DNA ligase in conjunction with splint oligonucleotides. Following this step, DNBs were generated and affixed to the custom-designed chips. After the sequencing phase, oligonucleotides containing a poly-T region and a 10-bp Molecular Identity (MID) were hybridized and ligated to the DNBs on the chips. The capture probes incorporated a 25 bp CID barcode, a 10 bp MID, and a 22 bp poly-T, which were primed for in situ capture. The CID sequences and their corresponding coordinates for each DNB were determined utilizing a base calling method, adhering to the manufacturer’s guidelines for the DNBSEQ sequencer. After sequencing, standard chips were separated from capture chips, ensuring that any duplicated CIDs in non-adjacent positions were eliminated.

### Stereo-seq tissue processing and imaging

The Stereo-seq experiments were performed according to our previously established protocols^11^. Initially, the Stereo-seq capture chip was rinsed with NF-H_2_O supplemented with 0.05 U/µL RNase inhibitor (NEB, M0314L) and subsequently air-dried at room temperature. Hippocampal tissue sections with 10 µL thickness were then applied to the chip surface and incubated at 37 °C for 3 minutes. Following this, the sections were fixed using methanol and stored at -20 °C for 40 minutes in preparation for the assembly of the Stereo-seq library. After fixation, the chips containing the tissue sections were stained with a nucleic acid dye (Thermo Fisher, Q10212), and imaging was conducted using a Motic Custom PA53 FS6 microscope in the FITC channel (10× objective) prior to the *in situ* capture.

### Stereo-seq *in situ* reverse transcription

Following the washing of the sections with 0.1× SSC wash buffer (Thermo, AM9770) containing 0.05 U/mL RNase inhibitor (NEB, M0314L), the tissue-coated chip was permeabilized using 0.1% pepsin (Sigma, P7000). This permeabilization involved incubation at 37 °C for 12 minutes, after which the sections were washed again with 0.1× SSC wash buffer containing 0.05 U/mL RNase inhibitor. The RNA released from the permeabilized tissue and captured by the DNB underwent overnight reverse transcription at 42 °C. This reaction utilized SuperScript II (Invitrogen, 18064-014) at a reverse transcriptase concentration of 10 U/µL, supplemented with 2.5 mM Stereo-seq-TSO (5-CTGCTGACGTACTGAGAGGC/rG//rG//iXNA_G/-3), 1× First-Strand buffer, 7.5 mM MgCl2, 5 mM DTT, 2 U/mL RNase inhibitor, 1 mM dNTPs, and 1 M betaine solution. Following reverse transcription, the tissue was digested using Tissue Removal buffer (25 mM EDTA, 10 mM Tris-HCl, 0.5% SDS, 100 mM NaCl) at 37 °C for 30 minutes. Subsequently, the chips containing cDNA were treated with Exonuclease I (NEB, M0293L) for 1 hour at 37 °C and were washed twice with 0.1× SSC.

### Amplification

The first-strand cDNAs obtained were amplified using KAPA HiFi Hotstart Ready Mix (Roche, KK2602) in conjunction with a cDNA-PCR primer (5’-CTGCTGACGTACTGAGAGGC-3’) at a concentration of 0.8 μM. The PCR amplification protocol was as follows: an initial denaturation at 95 °C for 5 minutes, followed by 15 cycles consisting of denaturation at 98 °C for 20 seconds, annealing at 58 °C for 20 seconds, and extension at 72 °C for 3 minutes, concluding with a final extension step at 72 °C for 5 minutes.

### Library construction and sequencing

The PCR products were purified utilizing VAHTS DNA Clean Beads (Vazyme, N411-03, 0.6×) and quantified with the Qubit^TM^ dsDNA Assay Kit (Thermo, Q32854). Subsequently, 20 ng of cDNA was fragmented using an in-house Tn5 transposase at 55 °C for 10 minutes. Upon completion of fragmentation, the reaction was halted by adding 0.02% SDS and gently mixing at room temperature for 5 minutes. The fragmented products were then subjected to the amplification phase. Specifically, 25 μL of the PCR product was combined with 1× KAPA HiFi Hotstart enzyme, along with 0.3 μM of Stereo-seq-Library-F primer (5’-CTGCTGACGTACTGAGAGGCA-3’) and 0.3 μM of Stereo-seq-Library-R primer (5’-GAGACGTTCTCGACTCAGCAGA-3’). The final reaction volume was adjusted to 100 μL by adding NF-H_2_O. The amplification protocol included one cycle at 95 °C for 5 minutes, followed by 13 cycles of 98 °C for 20 seconds, 58 °C for 20 seconds, and 72 °C for 30 seconds, concluding with a final extension at 72 °C for 5 minutes. Following amplification, the products were purified with AMPure XP Beads at purification volumes of 0.6× and 0.15× the original reaction volume. The purified products were then utilized to generate DNBs. Finally, these prepared DNBs underwent sequencing using the MGI DNBSEQ-T10 sequencer at the CNGB.

### Stereo-seq data Bin100 processing

The spatial transcriptomic gene expression profile matrix was segmented into non-overlapping bins, each covering an area of 100 × 100 DNB (bin100)^6,12^. Unique Molecular Identifiers (UMIs) were aggregated per gene within each bin, resulting in a bin100-gene matrix for subsequent analysis.

### Image-based segmentation of single cells in Stereo-seq data

Cell segmentation of Stereo-seq data was performed by StereoCell which was previously reported^26^. Briefly, first, we calculated the total UMI counts on the corresponding spatial coordinates of each DNB point and constructed a spatial density matrix. This matrix was then transformed into an image representation, where each pixel represents a DNB point and the intensity of the pixel’s gray reflects the UMI total number. Next, we aligned the nucleic acid staining images with Stereo-chip data from the same tissue slices by manually registering them. Cell segmentation was used based on ESPANet^98^ (https://arxiv.org/abs/2105.14447) optimization and training models, aims to enhance the capacity and long-distance multi-scale said channel dependence. After the segmentation, the watershed algorithm was used for post-processing, and the black and white mask of the cell was generated. For each segmented cell, we pooled UMI counts from all DNBs and aggregated them by gene, forming a cell-gene matrix for subsequent analysis.

### Cell type annotation of segmented cells

The annotation of segmented cells in Stereo-seq data was performed by our previously published method Spatial-ID^27^. A four-layer deep neural network (DNN) was initially trained on single-nucleus RNA sequencing (snRNA-seq) data, utilizing common genes as anchors to generate cell type probabilities for each cell. We constructed an adjacency matrix to encode spatial neighborhood information based on normalized Euclidean distances. A graph convolution network (GCN), comprising deep autoencoders and a classifier, was then applied. The autoencoders processed both the adjacency matrix and gene expression profiles, employing dual reconstruction losses via self-supervised learning. The classifier aligned these representations with initial probabilities through supervised learning. After 200 epochs of optimization, cell types were assigned based on the highest GCN-generated probabilities. By effectively aligning cell types from snRNA-seq data, Spatial-ID reduced batch effects in the cell type annotation process across Stereo-seq sections.

### Correlation analysis of cell types between Stereo-seq and snRNA-seq data

The snRNA-seq and Stereo-seq datasets were normalized using the Seurat package in R. This process included the selection of the high variable genes for subsequent analysis. Spearman correlation coefficients were then calculated using the overlapping top variable genes identified in both the snRNA-seq and Stereo-seq datasets. This approach aimed to evaluate the correspondence and relationship between cell types across the two datasets.

### Neighborhood complexity analysis in Stereo-seq data

Neighborhood complexity analysis was conducted using Squidpy (v1.3.0) as described by Palla et al. (2022)^99^. The connectivity matrix was initially computed for each individual section and subsequently aggregated by calculating the mean scores. Cell type cluster proximity was assessed using a permutation-based test (1,000 permutations), which involved comparing the actual cell type labels to a randomized configuration while preserving positional information. For each pair (actual label versus each permutation label), the mean and standard deviation were estimated, followed by the calculation of a Z-score to quantify the degree of cell type clustering.

### Cellbin trajectory construction

The clean counts matrix of cell type was imported into the R package SingleCellExperiment. A function slingshot was used to construct the trajectory by setting the parameter start.clus to indicate the starting point and not specifying the end point, and finally obtain the trajectory of this cell type. Subsequently, the clean counts matrix of the trajectories predicted by slingshot were input into the R package monocle3^38^. The functions reduceDimension and orderCells were used to construct the trajectory. Then, according to the pseudotime of the trajectory predicted by slingshot^100^, the differentialGeneTest function was used to calculate the genes with pseudotime changes. Finally, the plot_pseudotime_heatmap function was used to visualize the pseudotime gene expression heatmap, and plot_genes_in_pseudotime was used to show the expression of each gene along the pseudotime.

### The surrounding cell type composition of a certain target cell type

Taking the target cell type as the center, we extended the radius to a distance of 50 µm, covering the ranges of 20 µm, 30 µm, 40 µm, and 50 µm in length and forming 4 cocentric cricles. Next, we calculated the number of cells belonging to each cell type in different intervals of cocentric circles, taking this as the composition of surrounding cells of the target cell type. When calculating the difference in the proportion of clusters in the sclerosis (-) and the sclerosis (+) groups, the proportion of each cluster around the target cell type of the sclerosis (+) group calculated by the above method was subtracted from the proportion of the sclerosis (-) group to obtain the difference.

### Cell type density calculation

For each chip, we divided the number of each cell type in each brain region by the number of bin100 or the area of each brain region to obtain the brain region density of the cell type in this chip. Then, for the same cell type, we calculated the average of its brain region density in multiple chips to obtain the final brain region density value of this cell type.

### Identification of genes specifically expressed in different DG subregions

We used the Findallmarkers function to individually calculate the specifically expressed genes within the DG region, specifically in MLDG, GLDG, and PLDG, for each chip. Genes that exhibited a log_2_FC > 0.15, *p* < 0.05, and appeared in more than 50% of the chips were retained as significantly differentially expressed marker genes.

### Identification of differentially expressed genes between EX_CAs and EX_DG in sclerosis (-)

We used the FindMarkers function to calculate the DEGs between EX_CA1 and EX_DG, as well as between EX_CA2-4 and EX_DG. Genes that exhibited significant differential expression in both EX_CA1 and EX_CA2-4, as well as those that were significantly differentially expressed in both calculations for EX_DG were retained.

### Identification of DG-specific expression genes in the sclerosis (-) group that were co-expressed in both DG and CA4 brain regions in the sclerosis (+) group

First, we calculated the genes specifically expressed in the DG brain region compared to other brain regions in each chip of the sclerosis (-) group using the FindMarkers function. Genes with a log_2_FC > 0.15, *p* < 0.05, and appearing in more than 50% of the chips were retained as significantly differentially expressed marker genes. We applied the same method to calculate the genes specifically expressed in the DG brain region relative to the CA4 brain region in the sclerosis (-) group. By intersecting the above two results, we obtained the genes that were specifically expressed in the DG brain region of the sclerosis (-) group but not in the CA4 brain region. Then, using the same methodology, we calculated the genes specifically expressed in the DG and CA4 brain regions compared to other brain regions in the sclerosis (+) group. We then intersected the genes specifically expressed in the DG brain region but not in the CA4 brain region of the sclerosis (-) group with the genes co-expressed in the DG and CA4 brain regions of the sclerosis (+) group. This allowed us to identify the genes that were specifically expressed in the DG brain region of the sclerosis (-) group but not in the CA4 brain region, and were concurrently co-expressed in both DG and CA4 brain regions in the sclerosis (+) group.

### Identification of differentially expressed region-specific genes between hippocampal sclerosis (+) and sclerosis (-) groups

To identify the differential expression of region-specific genes between hippocampal sclerosis (+) and sclerosis (-) groups, we calculated the Pearson correlation coefficient of the logFC values of these specific genes, relative to other brain regions, between the two groups.

### Gene ontology analysis to determine the biological pathways

We first performed gene ontology analysis using “clusterProfiler” R package (v3.19), and then used the “rrvgo” R package (v3.19) to determine the biological pathways affected by cell type-specific DEGs by semantically clustered the significantly enriched terms into biological themes. Next, we set a cutoff of *p* < 0.05 and gene count > 4, then selected the most significant GO term from each biological theme to examine their enrichment levels in various cell types, represented by the *q* value.

### Spatial distance-constrained cell-cell communication analysis

We employed the StereoSiTE package to analyze cell-cell communication among the cell groups of interest in the spatial transcriptome data. Our analysis utilized 1,939 ligand-receptor (LR) interactions sourced from the CellChatDB human database, which categorizes interactions into three types: ‘Secreted Signaling’, ‘ECM-Receptor’ and ‘Cell-Cell Contact’. To assess communication probabilities between the selected cell groups, we utilized the intensities_count function with a distance threshold set to 50 µm. We filtered the ligand pairs based on a significance level of *p* < 0.05. The intensity of these inferred cell-cell communications was then visualized using the ggplot2 package, providing a comprehensive representation of the communication dynamics between the relevant cell groups.

### Tissue analysis and Single-nucleus suspension preparation

The methodology for isolating individual nuclei was implemented adhering to a previously established protocol^101,102^. In summary, the criteria for quality control in this study were met when the nucleus integrity was ≥ 85%, the clumping rate was ≤ 10%, and the total number of nuclei was ≥ 100,000. Four 100 μm frozen sections collected from each donor were underwent a thawing process, followed by mincing and transfer into a 1-ml Dounce homogenizer (sourced from TIANDZ). Additionally, 1 ml of homogenization buffer A was incorporated, comprising 250 mM sucrose (from Ambion), 10 mg/ml BSA (Ambion), 5 mM MgCl_2_ (Ambion), 0.12 units per ml of RNasin Plus (Promega, N2115), an equivalent concentration of RNase inhibitor (also Promega, N2115), and a comprehensive protease inhibitor cocktail (Roche, 11697498001) at 1× strength. The frozen tissues, kept chilled in an icebox, were homogenized using the loose pestle (designated as pestle 1) for a range of 25 to 50 strokes. This mixture was then passed through a 100-μm cell strainer into a 1.5 ml Eppendorf tube. Next, 750 ml of buffer A, now supplemented with 1% Igepal (Sigma, CA630), was added to a fresh 1-ml Dounce homogenizer. The tissue was further homogenized with the tight pestle (designated as pestle 2) for 25 strokes. Following filtration through a 40-μm strainer into another 1.5-ml tube, the mixture underwent centrifugation at 500 times gravity for 5 minutes at 4 degrees Celsius, resulting in the formation of a pellet containing nuclei. The obtained pellet was resuspended in 1 ml of buffer B, specifically tailored to contain 320 mM sucrose, 10 mg/ml BSA, 3 mM CaCl_2_, 2 mM magnesium acetate, 0.1 mM EDTA, 10 mM Tris-HCl, 1 mM DTT, the same protease inhibitor cocktail at 1× strength, and 0.12 units/ml of RNase inhibitor. The resuspended nuclei were centrifuged once again under the same conditions, yielding another nuclei-rich pellet. Finally, these nuclei were resuspended in a cell resuspension buffer to achieve a concentration of 1,000 nuclei/μl, ready for subsequent library preparation steps.

### snRNA-seq library construction and sequencing

Employing the DNBelab C Series Single-Cell Library Prep Kit (MGI, ID 1000021082), we generated barcoded libraries from single-nucleus suspensions^103^. This involved several key steps: droplet formation, emulsion disruption, bead harvesting, reverse transcription, and finally, cDNA amplification. The indexed libraries were then crafted in accordance with the manufacturer’s specifications, and their concentrations were accurately assessed using the Qubit ssDNA Assay Kit (Thermo Fisher Scientific, Q10212). Subsequently, the libraries underwent sequencing on the DNBSEQ-Tx platform at the China National GeneBank located in Shenzhen, China. The sequencing strategy employed featured distinct read lengths, with read 1 set at 41 base pairs and read 2 extended to 100 base pairs, offering a balanced approach for comprehensive genomic analysis.

### snRNA-seq data processing

The raw sequencing data acquired from DNBSEQ-TX underwent a process of alignment, specifically targeting the GRCh38.p12 human genome reference. Notably, pre-mRNA encompasses transcripts that were yet to undergo splicing to exclude introns. By incorporating intron readings from these pre-mRNA molecules into the ultimate gene expression tally, we ensured that no information is overlooked from the pre-mRNA stage. As a result, the transcript count for each gene in the snRNA-seq analysis was comprehensively determined, including both exon and intron reads. The ambient RNA noise was primarily from non-cell-specific mRNA contamination in the input solution. This contamination arose from the release of mRNA from ruptured cells into the solution. The ambient RNA noise could hinder accurate cell type identification, interfere with the analysis of gene expression differences, and obscure the true signals of low-expressed genes. Consequently, SoupX (v1.6.2; https://github.com/constantAmateur/SoupX) was employed here to mitigate the ambient RNA noise^104^.

### Quality control and cell clustering of snRNA-seq dataset

The quality control and clustering analysis of the snRNA-seq dataset was performed using Seurat (v4.2.2) and the R program. In specifics, we used “DoubletFinder” (v2.0.3) to locate and eliminate the doublets. Prior to further use, all query genes were guaranteed to be expressed in at least three cells. Low-quality cells were defined as those with fewer than 500 detected genes or more than 98% of cells in gene number in each library. Additionally, cells with a mitochondrial gene ratio of more than 10% in data preprocessing were filtered out. The “FindVariableFeatures” program was used to choose the top 3000 highly variable genes. Principal component analysis (PCA)-based dimension reduction was first carried out for downstream clustering and visualization. Then, the first 30 principal components (PCs) were retrieved for a subsequent Louvain clustering to identify the cell clusters (the resolution was set to 0.5). Finally, the clustering results were characterized in a two-dimensional space using the UMAP. Then, using the FindAllMarkers function with the following parameters: *p*.adjust < 0.05, avg_log_2_FC > 0.25, min.pct = 0.1, the cell clusters were annotated using known biomarkers that were more highly expressed in a given cluster. Significance at *p*.adjust < 0.05 was determined using adjusted *p*-values (Wilcoxon test).

### Multiple-sample integration and batch correction

To conduct cross-sample analysis, we combined data derived from six distinct samples. Ensuring the accuracy of the experiments as well as the analysis, two libraries were included for each sample. Prior to the integration process, each library’s data underwent meticulous preprocessing, encompassing stringent quality assurance measures, elimination of suboptimal cells, and data normalization to ensure consistency. Then, we merged the preprocessed data using the merge function and then performed linear dimensionality reduction analysis through PCA. The NormalizeData in the Seurat package was used to normalize the expression matrix of each cell after filtering. The “FindVariableFeatures” function was used to identify the top 3000 highly variable genes. The ScaleData in the Seurat package was then used to linearly scale the single-cell RNA-seq data. Batch corrections were performed for different samples using the Harmony software package (https://github.com/immunogenomics/harmony)^105^.

### CA1 region digitization and gradient gene clustering

The Spateo package^106^ was capable of digitizing the columns of a spatial domain by solving partial differential equations, where spateo was used to divide the proximal and distal ends of CA1 into three groups each. After obtaining six digitized brain regions, the moanin function was used to identify the spatially graded genes differentially expressed in sclerosis (-) and sclerosis (+). The splines_kmeans function performed k-means clustering on the gene expression data with the specified parameter n_clusters = 6. The splines_kmeans_score_and_label function evaluated and labeled the fit of all genes to the cluster centers. After obtaining the clustered genes, GO enrichment of genesets was performed to annotate the primary function of each geneset.

### Pseudospatial reconstruction of single-cell transcriptomes

Trajectories were constructed according to the procedure recommended in the Monocle2^39^ documentation (http://cole-trapnell-lab.github.io/monocle-release/docs/#constructing-single-cell-trajectories). Briefly, genes used to order cells were selected by comparing the inner and outer cell fractions in the assay. For each sample of EX_CA1 cells, differential gene expression analysis was performed between differentialGeneTest function in Monocle2. The cells were reduced dimensionality by DDRTree method. Based on the plot_cell_trajectory function, the cells were sorted and visualized, and colored by time and CA1 subregions, respectively. Expression of key markers across pseudospace was visualized using the plot_genes_in_pseudotime function in Monocle2 specifying a minimum value of 0.1 (min_expr = 0.1).

### Subregion parcellation of hippocampus

H&E staining images, spatial transcriptome data from neighboring slices, and ssDNA and totalRNA images from transcriptome data microarrays were combined to precisely delineate brain regions. Initially, normalized SingleCellExperiment objects were created using the transcriptome data of the defined partitions (bin100) on each chip as input for the ensuing BayesSpace clustering^107^. The first 50 principal components of the first 3,000 highly variable genes were kept after PCA was conducted on them. With the help of the bayes clustering subgroup marker genes that were retrieved via the FindAllMarkers function, brain areas were annotated and defined for outlines. With the help of the image processing R package (imager: http://dahtah.github.io/imager/), the defined 2D brain area outlines were recognized. The contours were refined in accordance with the cell morphology, density, and staining intensity in these photos after being combined with H&E staining images, ssDNA images, and totalRNA images, respectively. The ultimate hippocampus subregion delineation was achieved.

### Dimension reduction and clustering

PCA and uniform manifold approximation and projection (UMAP) was used for the dimensionality reduction analysis. The FindNeighbors and FindClusters in the Seurat package were used for cell cluster analysis. Through the FindAllMarkers in the Seurat package, marker genes for each cell cluster according to the following screening criteria: *p*.adjust < 0.05, |log_2_FC| > 0.25 and min.pct = 0.1.

### Analysis of differences within brain regions as well as cell types

For differentially expressed genes between sclerosis (-) and sclerosis (+) in different cell types as well as in different brain regions, the list of differentially expressed genes between cell types was calculated using the FindMarkers function in Seurat and filtered with the following settings (min.pct = 0.1, logfc.threshold = 0.25, only.pos = TRUE).

### Gene set analysis for Stereo-seq bin100 data

Following the enrichment of relevant pathways using DEGs, we scored the gene sets enriched in these pathways, specifically in both the sclerosis (-) and sclerosis (+) groups, utilizing the AUCell package (https://github.com/aertslab/AUCell). The resulting scores were then compared by subtracting the scores of the sclerosis (-) group from the sclerosis (+) group, allowing for further validation of the enrichment status of these pathways.

### Immunohistochemistry

Three biological replicates were performed for each immunohistochemistry experiment. Briefly, hippocampal sections were incubated at room temperature with 0.3% HLOL for 30 minutes to block endogenous peroxidase activity. After rinsing with PBS, sections were treated with 0.3% Triton X-100 for 10 minutes to permeabilize the tissue. Sagittal sections were then incubated with horse serum at room temperature for 1 hour to block nonspecific binding. The hippocampal slices were incubated overnight at 4°C with the NeuN primary antibody (1:500, Abcam, ab177487) and the GFAP primary antibody (1:2500, Proteintech. Cat #16825-1-AP) respectively. After three PBS washes, sections were incubated with a biotinylated secondary antibody and Vectastain ABC solution (Vector Labs, Burlingame, CA) for 1 hour at room temperature. Slides were mounted with ProLong Gold Antifade Mounting Medium (Thermo Fisher, Cat#P-36931) and sealed with 2.0 mm² coverslips using clear nail polish. Negative controls (no primary antibody) were included in each immunohistochemistry experiment. Hippocampal sections were examined under a light microscope. For each patient-derived sample, three sections were analyzed at 100 μm intervals.

### Calculation of cell density in immunostaining

We employed FiJi (https://imagej.net/software/fiji/downloads) for quantitative density calculations in the hippocampus. The percentage of NeuN positive cells was determined manually using FiJi Cell Counter plug-in. The area of hippocampal subregions was quantified using the “Area Measure” tool in Fiji. Cell density was calculated by dividing the number of NeuN-positive cells by the area of each hippocampal subregion.

### Chromogenic Substrate and Image Capture

DAB chromogenic substrate solution, 100 μL per section, was applied, and the color development was controlled under a microscope for 3-5 minutes; if no color was visible after 15 minutes, it was considered negative. The color development was terminated with a PBS buffer. The sections were then counterstained with hematoxylin for 1 minute, dehydrated through a gradient ethanol series for 5 minutes, cleared in xylene for 3 minutes, and mounted with neutral resin. All operations were performed while keeping the tissue moist; all added reagents completely covered the tissue. The sections were scanned and the results were analyzed.

### Statistical Analysis Methods

Data were analyzed using appropriate statistical methods based on the type of variable. For categorical variables, Yates’ correction was applied to account for small sample sizes in contingency tables. Continuous variables were tested for differences among groups using one-way analysis of variance (ANOVA). Post-hoc comparisons between groups were performed using Tukey’s Honest Significant Difference (HSD) test. Kruskal-Wallis test was used for comparing data of multiple groups, the Spearman correlation coefficient test was used for correlation analysis of signal pathway activity strength in different directions with different gradient distances; linear regression and Pearson correlation tests were used for correlation analysis of gene expression levels with signal pathway activity. Data were tested bilaterally, and a *p* < 0.05 was considered statistically significant.

## Acknowledgments

This work was supported by National Key R&D Program of China (2022YFC2503800), National Science Foundation of China (82471388, 31900397), the National Science and Technology Innovation 2030 Major Program (2021ZD0204400), Clinical Research Incubation Project of West China Hospital of Sichuan University (22HXFH022), Joint Fund of the National Natural Science Foundation of China (U21A20393), 1.3.5 project for disciplines of excellence from West China Hospital of Sichuan University (ZYGD23032) and Science & Technology Department (No.2021YFC2401204, 2022YFC2503805). We thank BGI for hosting various datasets from this study. We thank CNGB (China National GeneBank) for hosting various datasets from this study. We would like to thank DCS Cloud (https://cloud.stomics.tech/) for providing the computational resources and software support necessary for this study.

## Author contributions

Z.H, L.H and LF.W conceptualized and designed the project; XJ.L collected tissue samples; XJ.L, YR.A, YN.X and LF.W designed the experiments; XJ.L, YR.A, YN.X performed the experiments; LF.W, YY.L, YR.W, BF.J and YZ.H performed and analyzed the sequencing data; LF.W, YY.L, YR.W, BF.J, YZ.H and XJ.L prepared the figures. XY.K, X.G, RX.Y, Y.M, R.T, JM.L, D.Z, L.P, Q.T, C.S, SP.L, L.L, X.X, and LQ.L provided relevant advices and reviewed the manuscript. LF.W, YY.L, YR.W, BF.J, YZ.H and XJ.L wrote the manuscript with input from all authors. All other authors contributed to the work. All authors read and approved the manuscript for submission.

## Additional information Supplementary information

Correspondence and requests for materials should be addressed to

---

**Extended Data Fig. 1.**
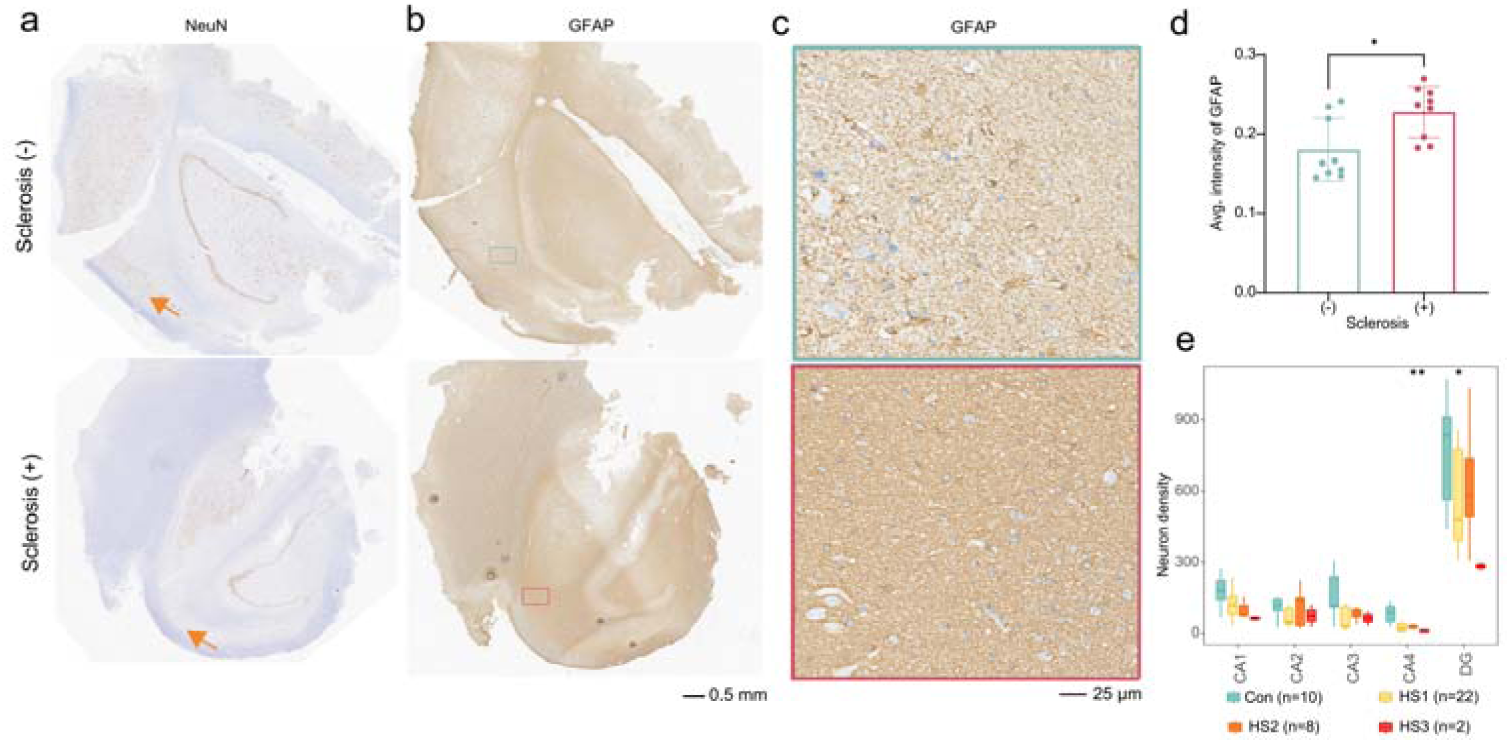
Diagnosis of hippocampal sclerosis. (**a**) NeuN staining of typical samples with HS (bottom) and without HS (top), arrow indicates the location where excitatory neuron lose in HS. (**b**) GFAP staining of typical samples with HS (bottom) and without HS (top), rectangle indicates the location where astrocyte was proliferated in sclerosis. (**c**) Zoom in the location marked in (b), border colors represent the origin of amplified area with (red, bottom) and without (green, top) sclerosis. (**d**) Bar plot indicates GFAP intensity of marked regions in sclerosis (-) and sclerosis (+), * *p <* 0.05, *t*-test. (**e**) Statistical quantification of NeuN staining for samples with and without sclerosis. Kruskal-Wallis test, ** *p* = 0.003.

**Extended data Fig. 2.**
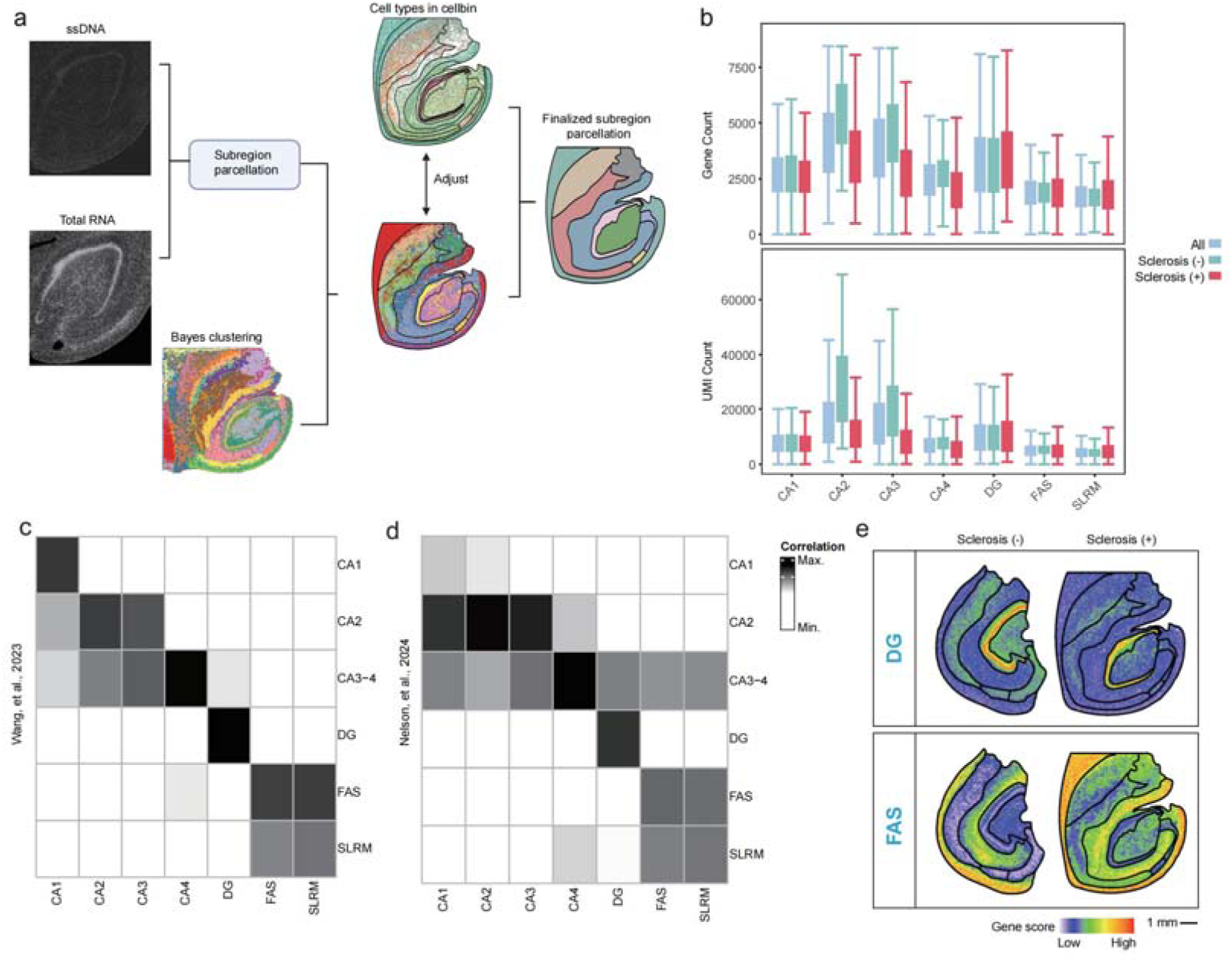
Confirmation of subregion parcellation for human hippocampus. (**a**) Schematic diagram of subregion parcellation for human hippocampal transcriptome. (**b**) Boxplot indicating the number of genes (top) and UMIs (bottom) per bin100 for each subregion. (**c-d**) Heatmap showing the correlation of parcellated subregions between Stereo-seq data and public dataset from Wang et al., 2023^14^ (**c**) and Nelson et al., 2024^14^ (**d**). (**e**) Spatial gene score profile of DEG module from **Fig.1c** between sclerosis (+) and sclerosis (-) in DG (top) and FAS (bottom).

**Extended Data Fig. 3.**
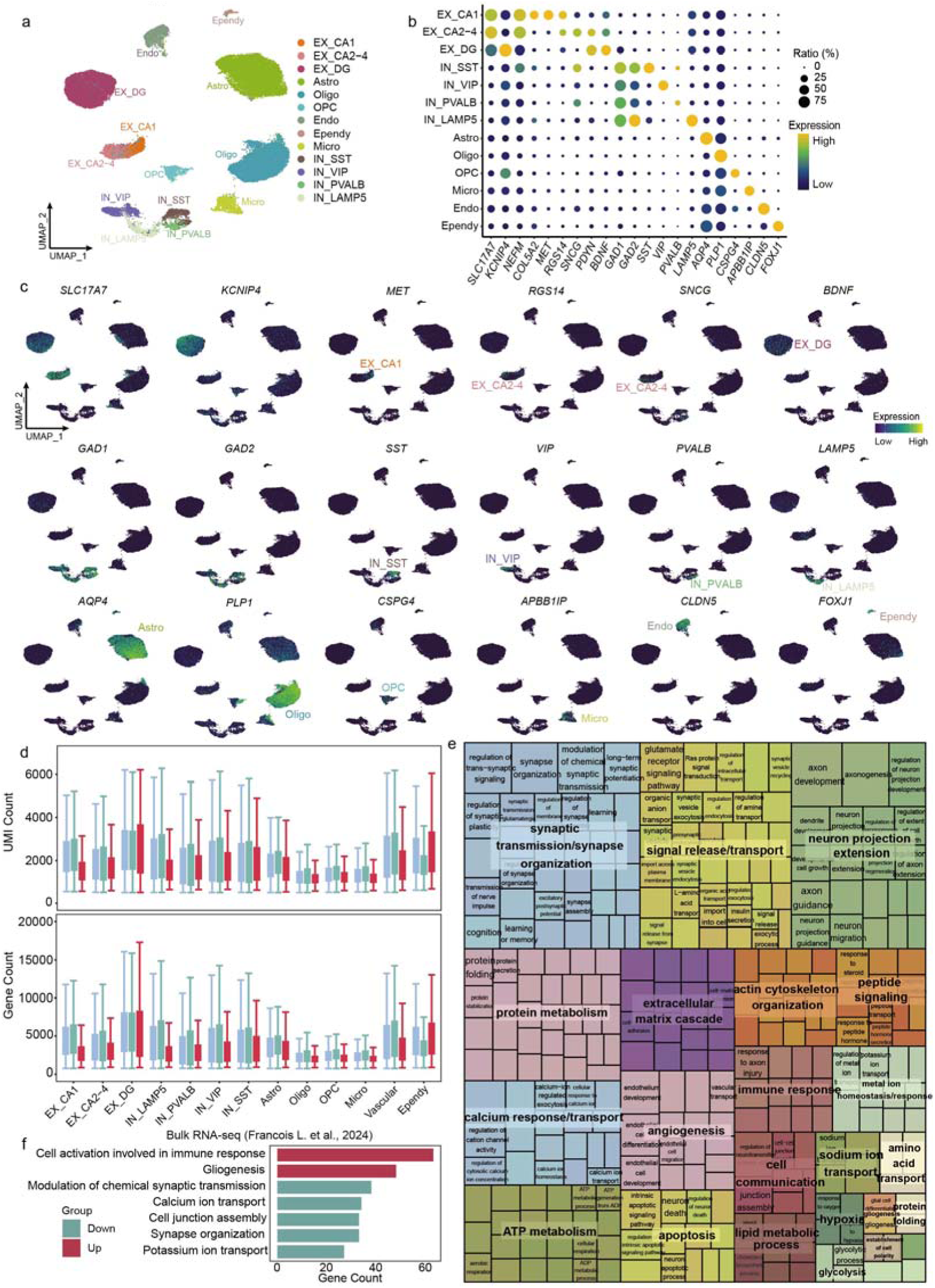
Cell type annotation and functional changes of snRNA seq. (**a**) Uniform manifold approximation and projection (UMAP) visualization of snRNA dataset. (**b**) Dotplot depicting the expression of marker genes in annotated cell clusters. (**c**) Expression of marker genes in UMAP. (**d**) Box plots showing the number of UMIs (bottom) and the number of genes (top) per cell in each cell type for total (Total), sclerosis (-), and sclerosis (+) samples in snRNA. (**e**) Aggregate enrichment of GO terms within each biological theme across cell types by down- and up-regulated DEGs, excluding themes implicated by only one term and nonsignificant cell types. Themes were named by the most significant term they contain. (**f**) Histogram showing functional enrichment of up-regulated (red) and down-regulated (blue) genes of public hippocampal sclerosis RNA dataset of human temporal lobe^8^.

**Extended Data Fig. 4.**
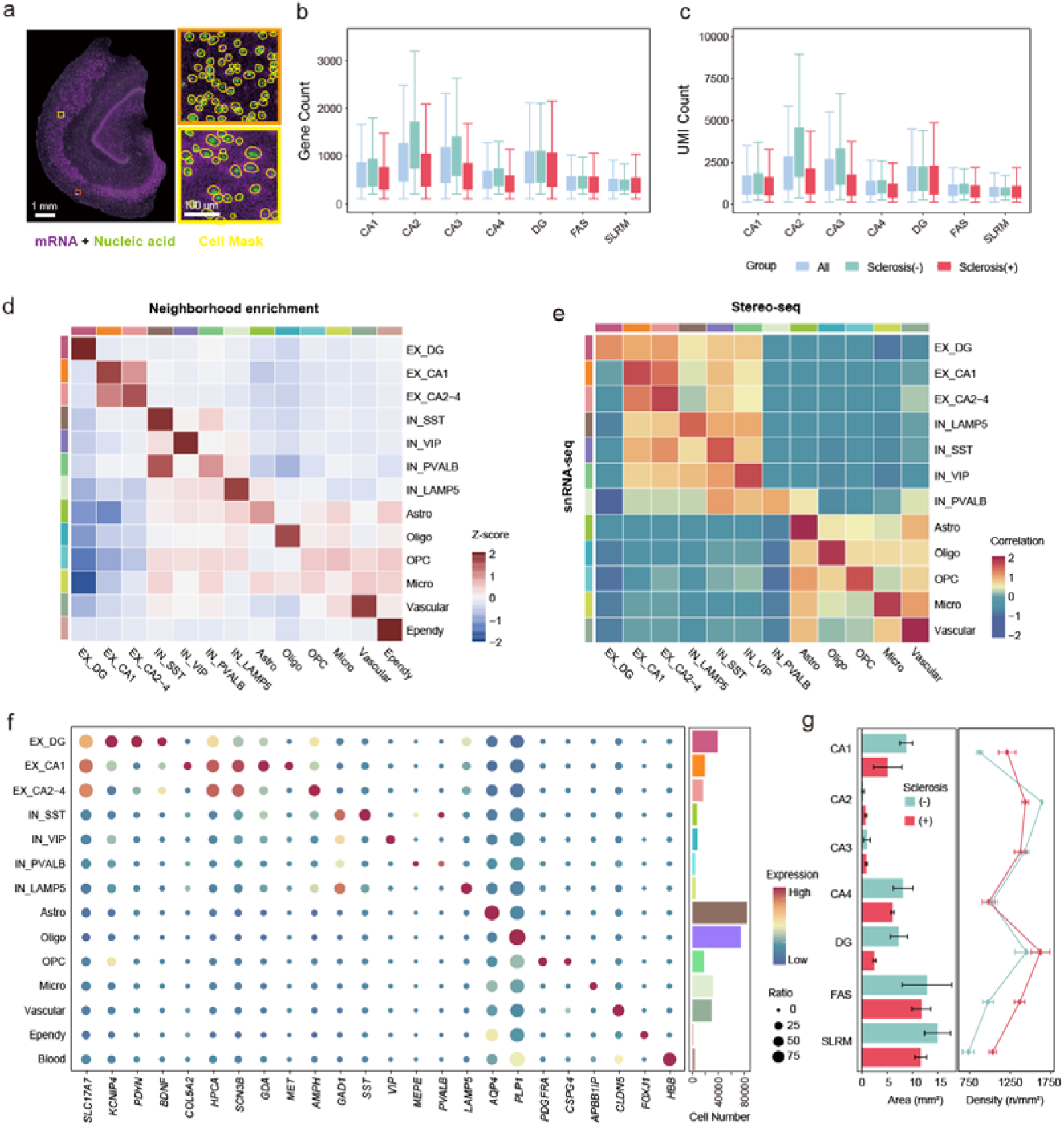
Single cell segmentation and annotation of Stereo-seq. (**a**) Single cell segmentation for Stereo-seq dataset. (left) AI-assisted automated segmentation based on superimposed nucleic acid staining (green) and total mRNA signals (purple) from a representative section of hippocampus sample. (right) Magnified areas were in yellow box of (left). Yellow circles represented boundaries of cells. (**b-c**) Box plots showing the number of genes (**b**) and UMIs (**c**) per cell in each subregion for all (All), sclerosis (-), and sclerosis (+) samples in segmented Stereo-seq. (**d**) Heatmap showing correlation between cell types annotated from snRNA-seq and Stereo-seq. (**e**) Heatmap showing neighborhood enrichment among segmented cell types annotated from Stereo-seq. (**f**) Dotplot showing expression profile of cell type-specific marker genes in spatial segmented and annotated cells. (**g**) Left: Bar plots showing the area in each subregion in both sclerosis (-) and sclerosis (+); right: cell density of each subregion in both sclerosis (-) and sclerosis (+).

**Extended Data Fig. 5.**
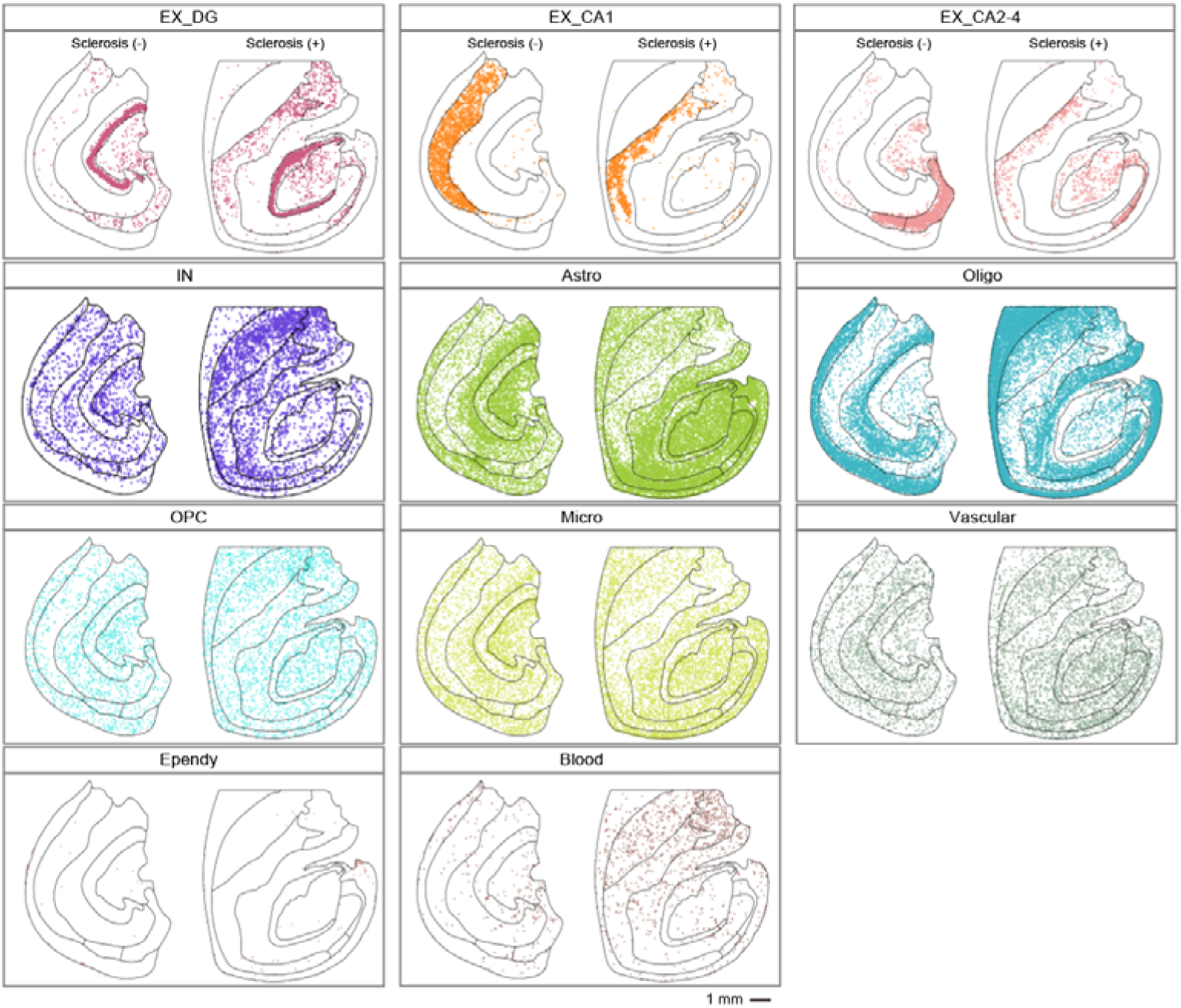
The spatial distribution of cell types with and without sclerosis. Spatial visualization of EX_DG, EX_CA1, EX_CA2-4, IN, Astro, Oligo, OPC, Micro, Vascular, Ependy and Blood cell.

**Extended Data Fig. 6.**
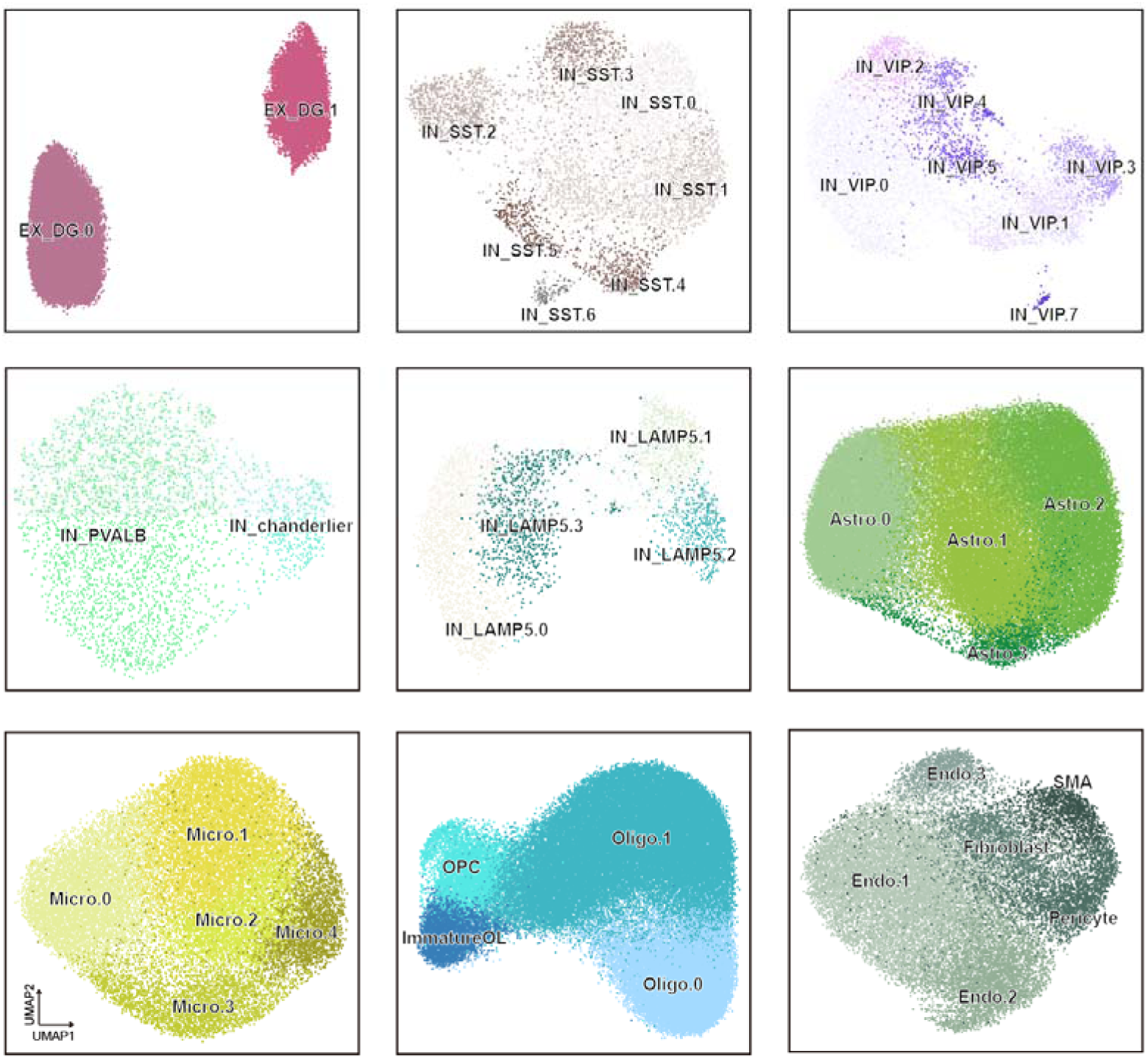
UMAP visualization of subtypes of 9 hippocampal cell types in Stereo-seq data. UMAP visualization of EX_DG, IN_SST, IN_VIP, IN_PVALB, IN_LAMP5, Astro, Micro, Oligo, OPC and Endo.

**Extended Data Fig. 7.**
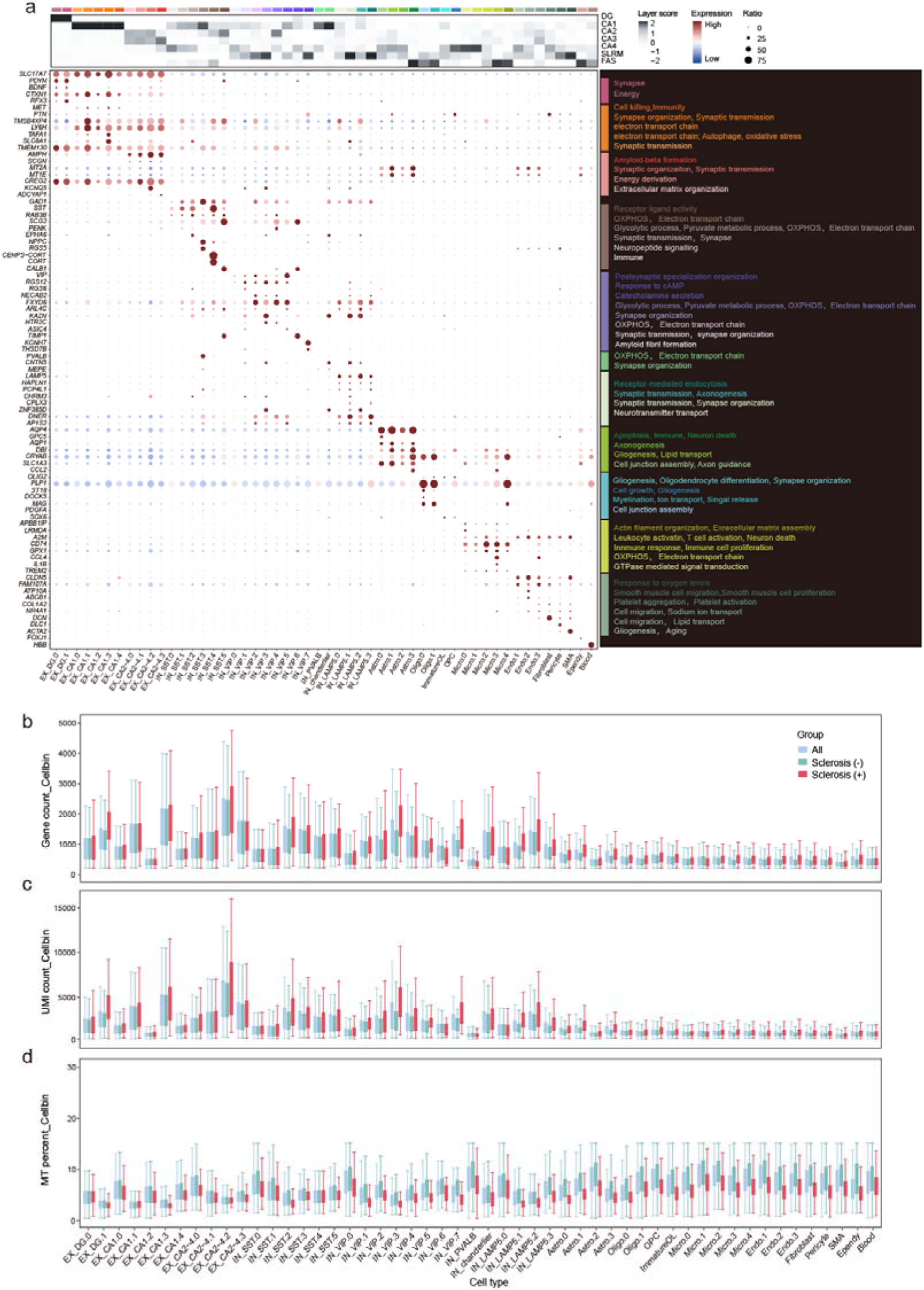
Annotation and quality of subtypes of hippocampus with and without sclerosis. (**a**) Dotplot showing the density change of each cell type in each subregion. The spot size represented average density of each cell type in each subregion over with and without sclerosis groups. The spot color represented an increase (red) or decrease (blue) of density in sclerosis. Comprehensive graph depicted characters of subtypes of hippocampus. (top) heatmap showing relative density of each subtype in each subregion, (bottom) dotplot showing marker gene expression of each cell subtype across the hippocampus, (right) words showing the enriched biological functions of each subtype. (**b-d**) Box plot indicating the number of genes (top), UMIs (middle) and MT percent (bottom) per subtype of all, sclerosis (+) and sclerosis (-).

**Extended Data Fig. 8.**
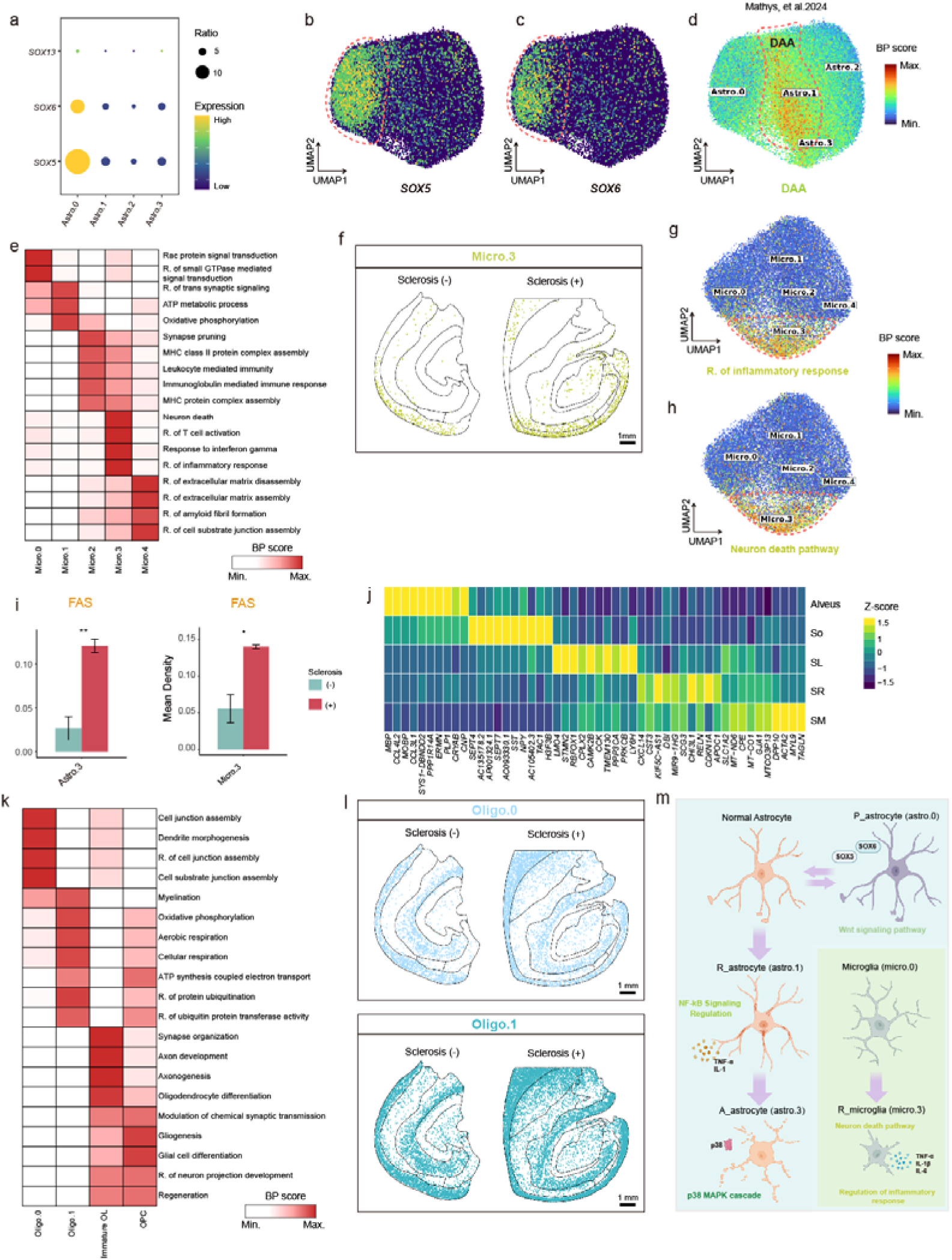
Distribution change of glial cell subtypes across subregions of hippocampus with sclerosis. (**a**) Dotplot showing the expression of *SOXD* family genes in astrocyte subtypes. (**b-c**) UMAP visualization of gene expression for *SOX5* (**b**) and *SOX6* (**c**) in astrocyte subtypes. (**d**) UMAP visualization of previous reported DAA related genes^30^ expression in astrocyte subtypes. (**e**) Heatmap showing gene expression change of selected enrichment pathways in microglial subtypes in sclerosis. (**f**) Spatial distribution of Micro.3 in hippocampal with (right) and without (left) sclerosis. Scale bar: 1 mm. (**g-h**) UMAP visualization of inflammatory response (**g**) and neuron death (**h**) related genes expression in microglia subtypes. (**i**) Bar plot showing cell density Astro.3 (left) and Micro.3 (right) in FAS with and without sclerosis. (**j**) Heatmap showing average expression of small area-specific genes in Alveus, SO, SL, SR and SM. (**k**) Heatmap showing expression change of selected enrichment pathways in oligodendrocyte and OPC subtypes. (**l**) Spatial distribution of Micro.3 in hippocampal with (right) and without (left) sclerosis. Scale bar: 1 mm. (**m**) Schematic diagram for subtypes of astrocyte and microglia.

**Extended Data Fig. 9.**
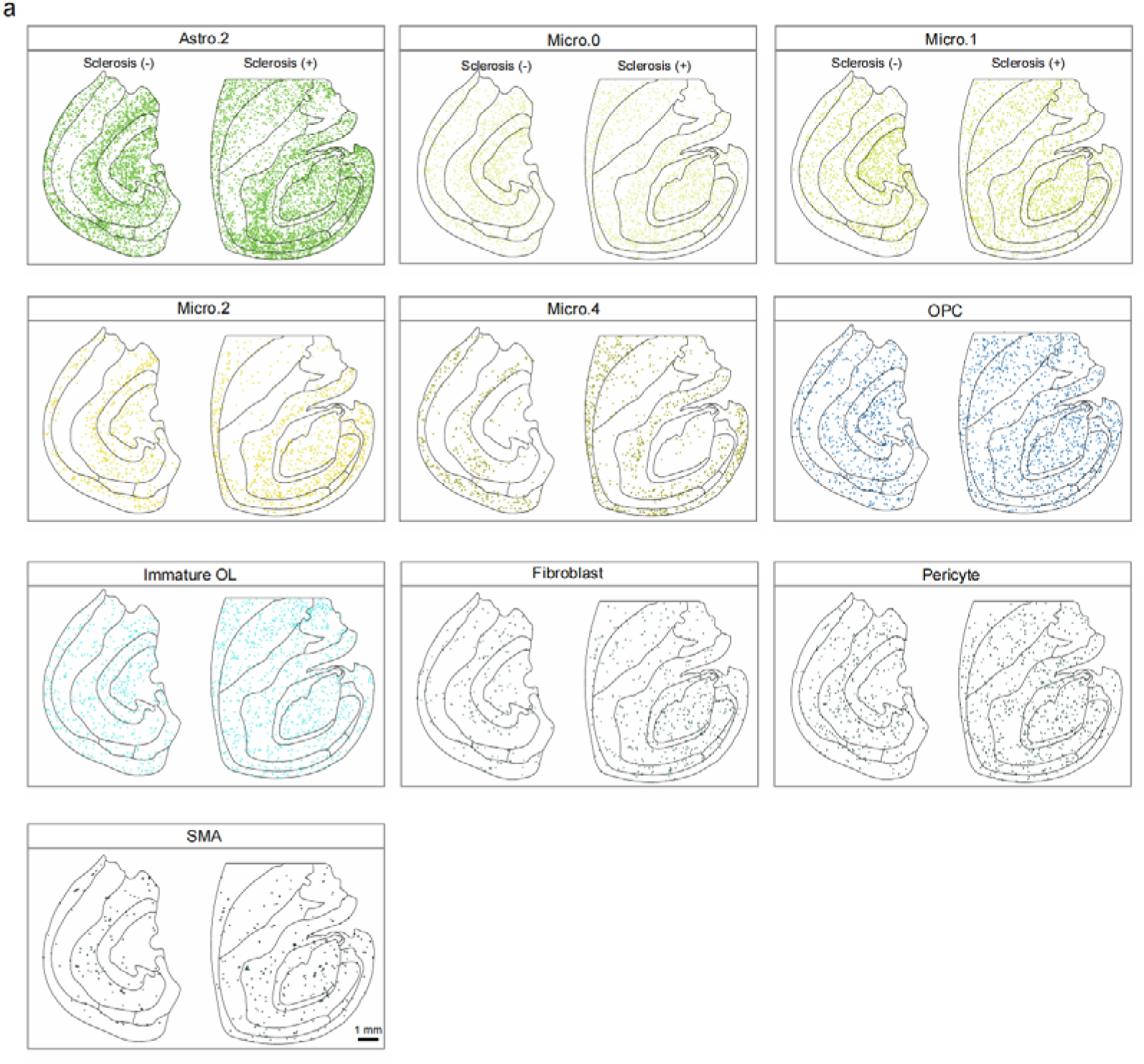
Spatial distribution of annotated subtypes of non-neurons from Stereo-seq data. Spatial visualization of subtypes of Astro, Micro, Oligo and Vascular cell. Scale bar: 1 mm.

**Extended Data Fig. 10.**
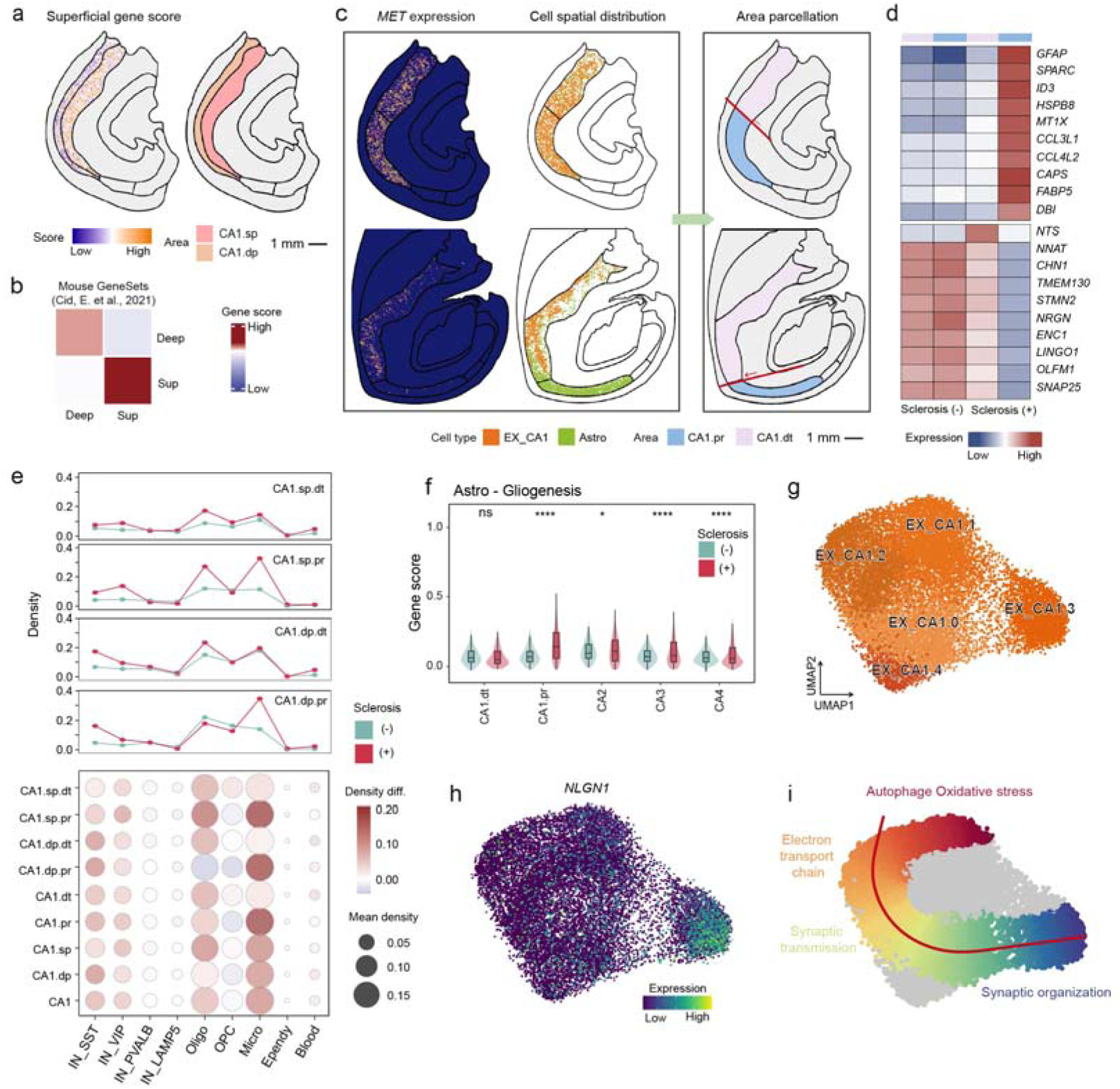
Cell type composition change of proximal and distal area in and surround CA1 with sclerosis. (**a**) The schematic diagram for CA1.sp and CA1.dp parcellation (right) based on spatial expression of CA1.dp marker genes^9^ (left). (**b**) Heatmap showing the average score of reported superficial and deep markers in mouse sclerosis model^9^. (**c**) The schematic diagram for proximal and distal CA1 parcellation (right) based on spatial expression of CA1 specific marker gene and spatial distribution of EX_CA1 and astrocyte (left). Scale bar: 1 mm. (**d**) Heatmap showing average expression of CA1.pr and CA1.dt specific genes of sclerosis in both sclerosis (+) and sclerosis (-). (**e**) Line chart (top) and dotplot (bottom) showing the density and density change of each cell type or subtype in each area of CA1 subregion. In dotplot, the spot size represented average density of each cell type in each subregion over with and without sclerosis groups. Density.Diff., Density difference. (**f**) Box plot revealing the gene expression score of gliogenesis in astrocyte in the CA1.dt, CA1.pr, and CA4 regions of sclerosis (+) and sclerosis (-) groups. (**g**) The UMAP of EX_CA1 subtypes. (**h**) The expression of *NLGN1* in EX_CA1 UMAP. (**i**) The UMAP plot of EX_CA1, colored by trajectory pseudotime (calculated by slingshot).

**Extended Data Fig. 11.**
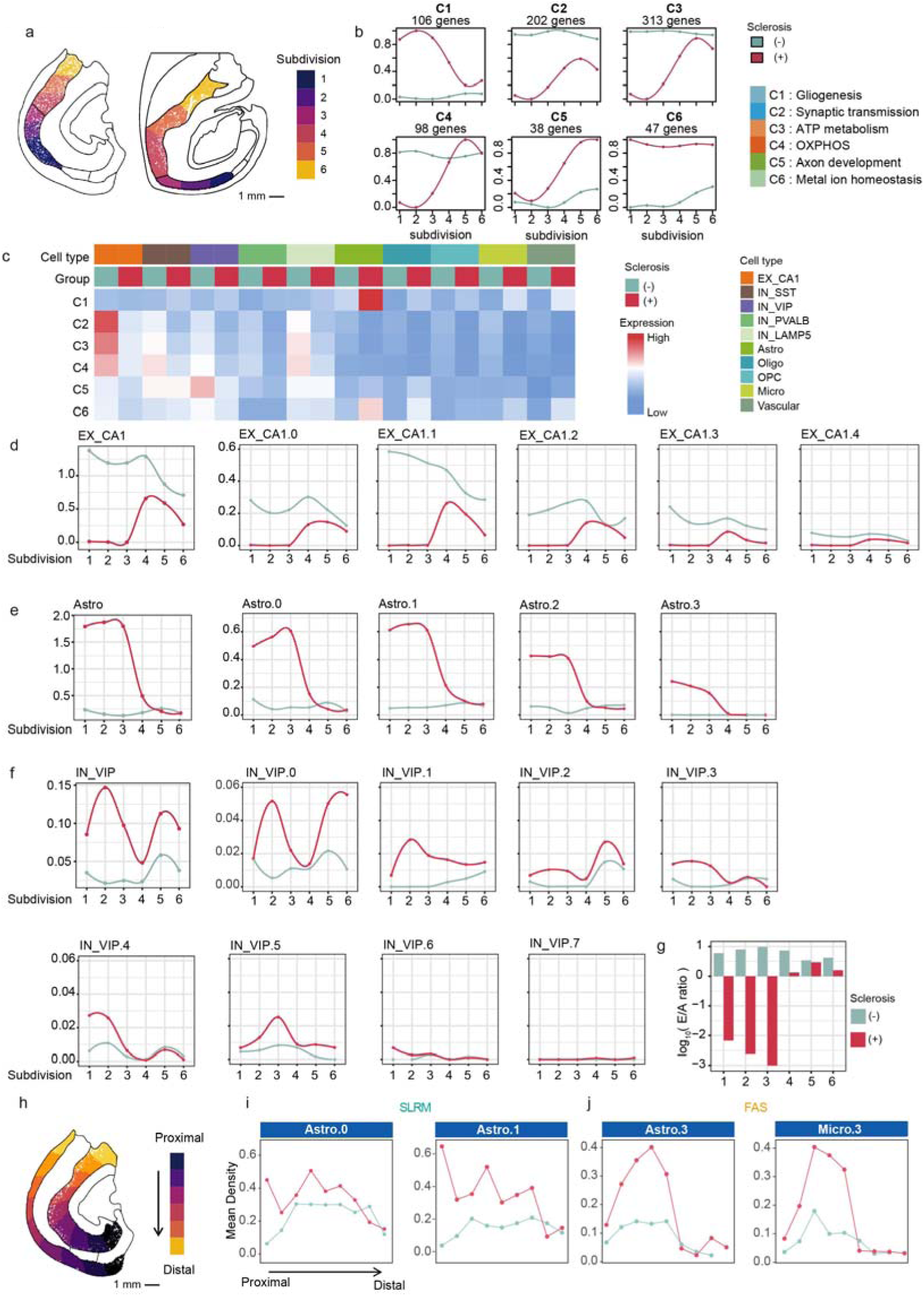
Categories of DEGs in sclerosis by each subdivision. (**a**) Digitized schematic diagram showing the CA1 proximal-distal axis for obtaining gradient expressed genes along each axis. (**b**) Line chart showing cluster of genes with different expression trends in CA1 proximal-distal axis with and without sclerosis. C1-C6 were six clusters with different expression trends and biological functions. (**c**) Heatmap showing the expression of each cluster in each cell type in CA1 subregion with and without sclerosis. The color represented higher (red) and lower (blue) expression in each cell type. (**d-f**) Density dynamics of cell type and subtypes for astrocyte (**d**), EX_CA1 (**e**), and IN_VIP (**f**) along with CA1 proximal-distal axis (CA1. p-d axis). (**g**) log_10_ (average E/A ratio) dynamic change with the digitization axis of CA1 E/A ratio: the density ration of excitatory neurons / astrocyte. The raw E/A density ratio was first adjusted by adding a minimal constant value (0.001) to avoid logarithmic singularity (i.e., handling cases where the denominator approaches zero). Subsequently, this adjusted ratio was converted to its logarithmic base 10 (log_10_) form. If the value is greater than 0, EX_CA1 density is higher than Astro, and vice versa. (**h**) Digitized schematic diagram showing the SLRM and FAS proximal-distal axis. Scale bar: 1 mm. (**i-j**) Density dynamics of subtypes for astrocyte and microglia along with SLRM (**i**) and FAS (**j**) proximal-distal axis.

**Extended Data Fig. 12.**
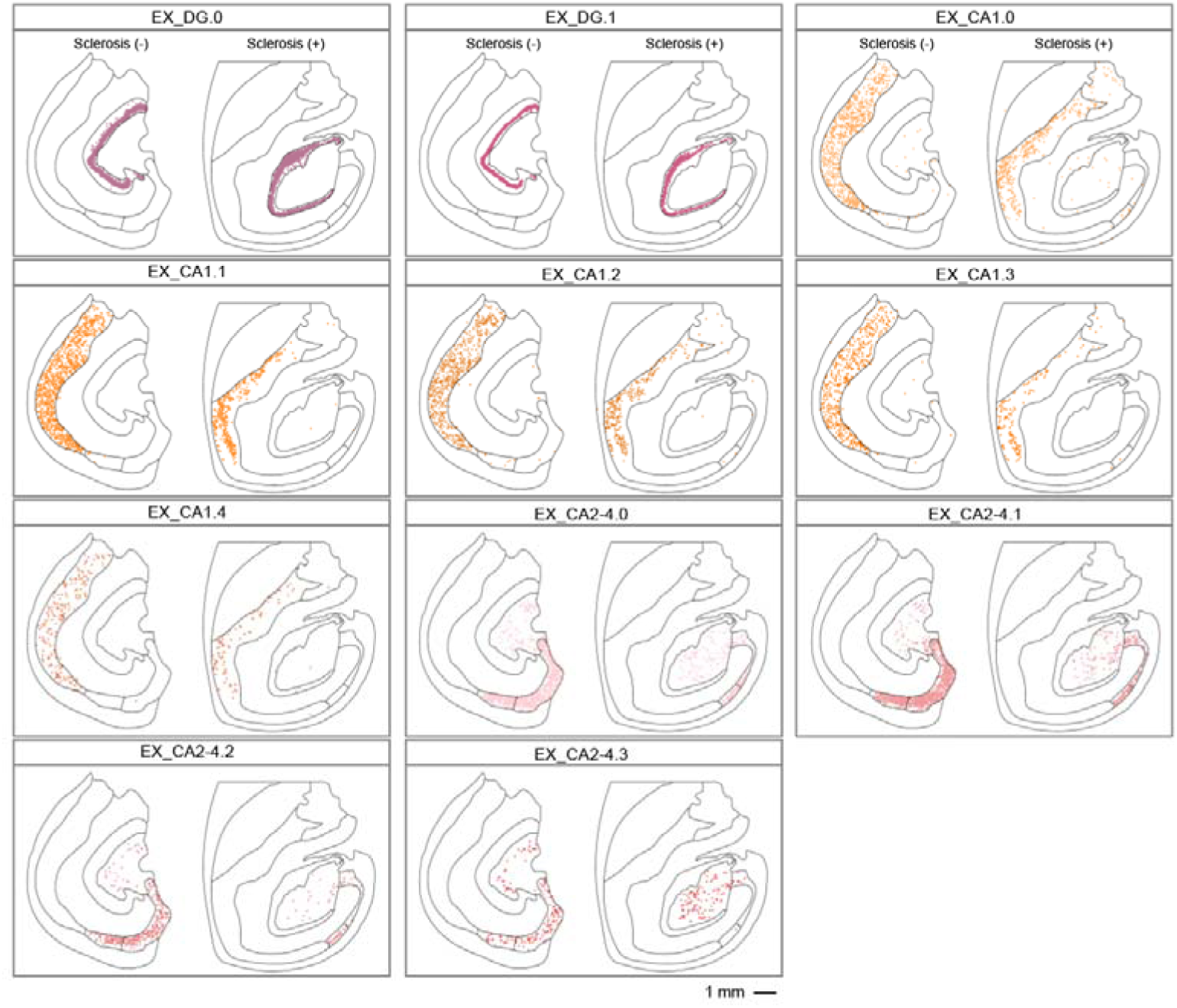
Spatial distribution of annotated subtypes of excitatory neurons. Spatial visualization of each subtype of EX_DG, EX_CA1 and EX_CA2-4. Scale bar: 1 mm.

**Extended Data Fig. 13.**
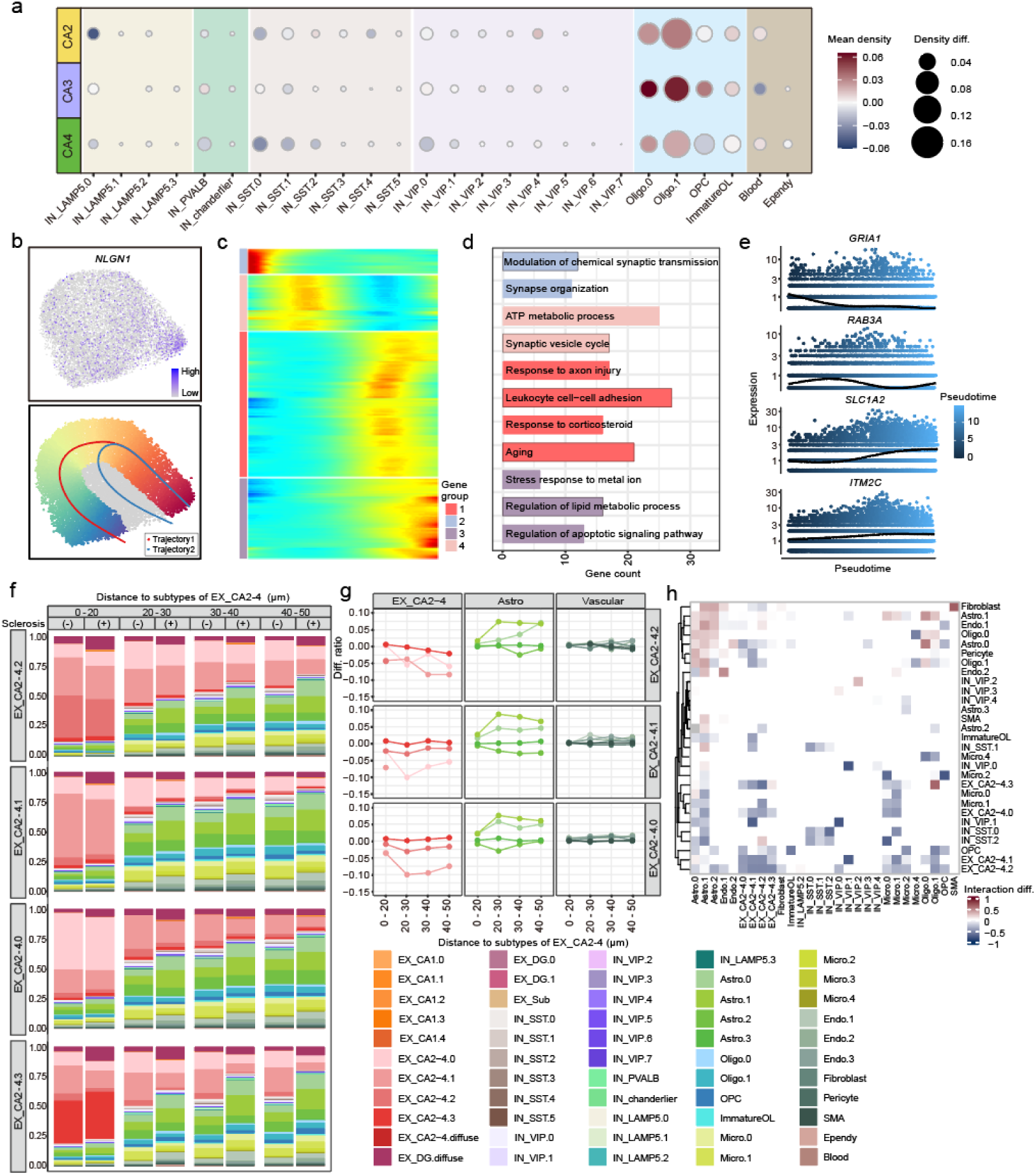
Pathological changes in the CA2-4 brain region affected by sclerosis. (**a**) Dotplot showing the density change of subtypes in each subregion. The spot size represented the average density of each cell type in each subregion of samples. The spot color represented increase (red) and decrease (blue) of cell density in sclerosis, respectively. Density. diff., Density difference. (**b**) Top: The expression of *NLGN1* in EX_CA2-4 UMAP. Bottom: UMAP colored by trajectory 1 pseudotime. (calculated by slingshot). (**c**) The heatmap displayed the expression of pseudotime genes along trajectory 1 and divided them into four groups based on the pattern of their gene expression along the pseudotime. Each row representing a single gene. (**d**) The bar chart displayed enriched GO functions of gene clusters in (**c**). The bars were filled with different cluster colors. (**e**) Kinetics plot showing the relative expression of representative genes in cluster 2 and 3 in (**c**) over pseudotime. Points were colored by trajectory 1 pseudotime. The height of the line represented the relative amount of gene expression. (**f**) The bar chart depicting the proportion of cell subtypes within various distances to each EX_CA2-4 subtype with and without sclerosis. (**g**) The line graph depicting the proportion difference of each subtype of EX_CA2-4 (left), astrocyte (middle) and vascular cell (right) between sclerosis (+) and sclerosis (-) groups at various distances to each EX_CA2-4 subtype in trajectory 1. (**h**) The heat map showed the difference in the interaction strength between different cell types with and without sclerosis. The colors represented increased (red) and decreased (blue) interaction in sclerosis.

**Extended Data Fig. 14.**
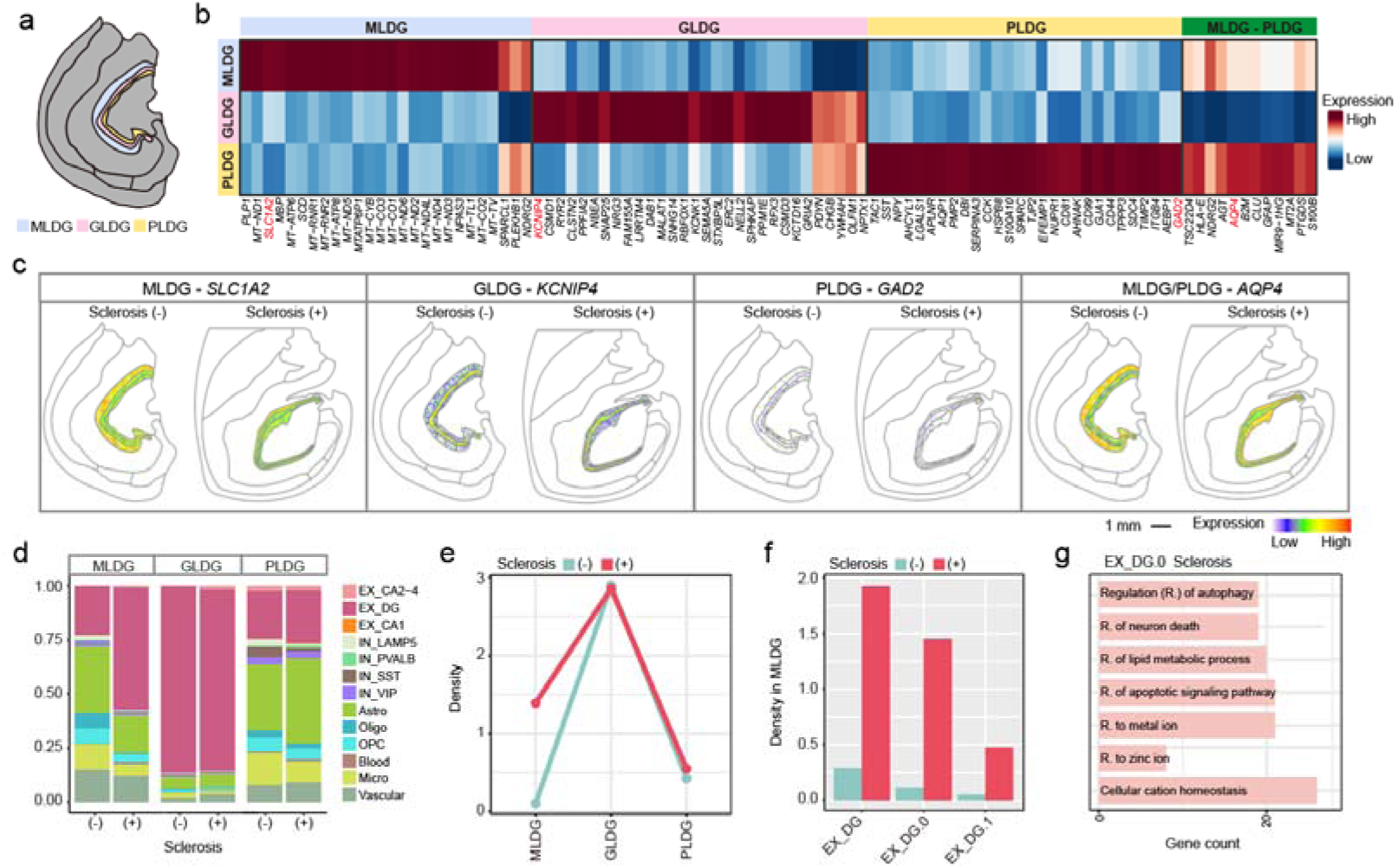
Pathological changes of DG subregions in HS. (**a**) Schematic diagram of DG region parcellation into three subregions. (**b**) The heatmap showing the expression of three area-specific genes of DG. (**c**) Spatial expression of representative genes in three subregions of DG. Scale bar: 1 mm. (**d**) The bar chart showing the proportion of subtypes in three areas of DG. (**e**) The line graph showed the density of EX_DG.0 in the three subregions of DG in sclerosis (+) and sclerosis (-) groups. (**f**) Bar plot showing the density of EX_DG and its subtypes in MLDG. (**g**) The bar chart showing the enriched functions of DEGs of EX_DG.0 in sclerosis.

**Extended Data Fig. 15.**
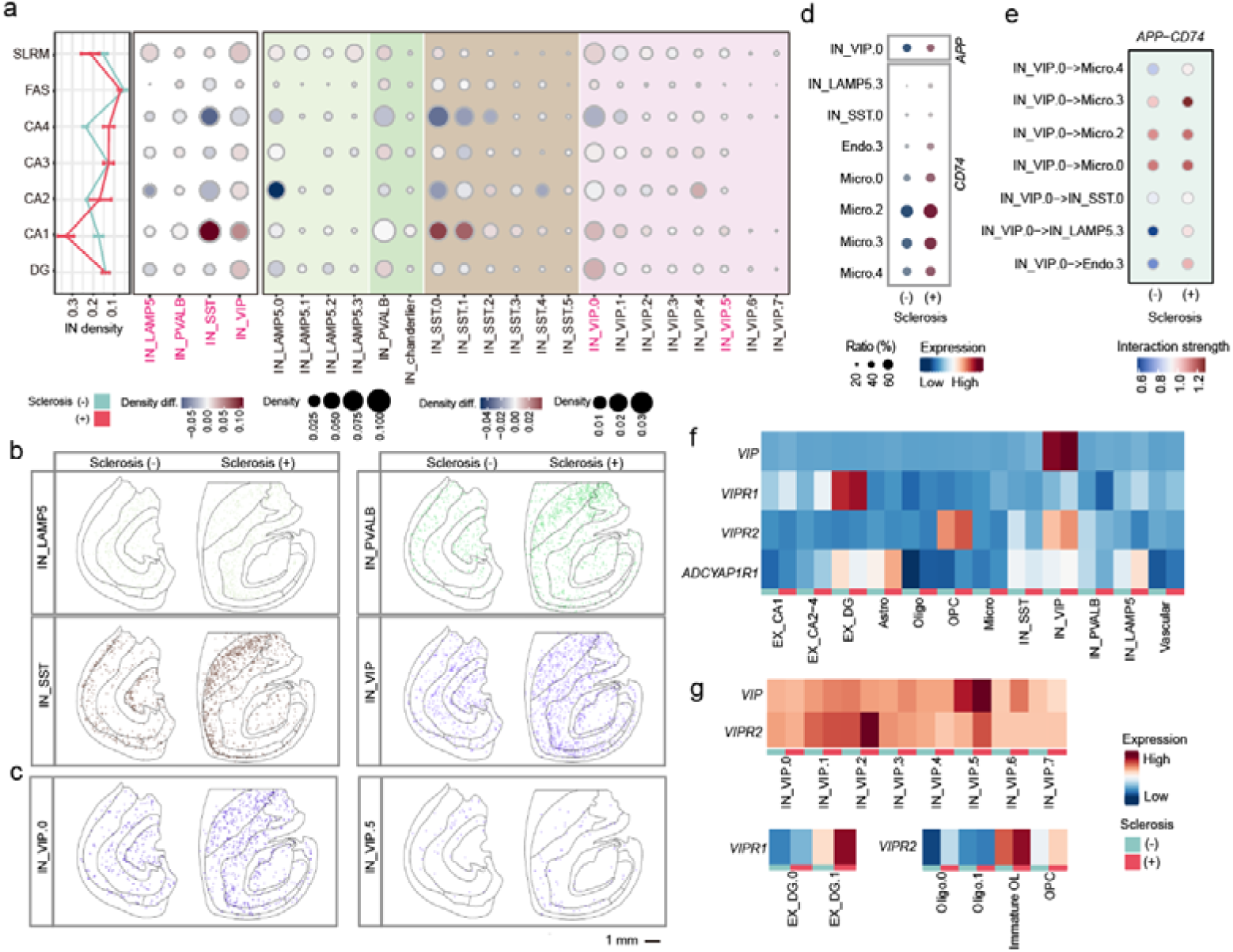
Distribution change of inhibitory neuron in sclerosis by each subregion. **(a)** Dotplot showing the density change of each cell type (left) or subtype (right) of inhibitory neuron in each subregion. The spot size represented average density of each cell type or subtype in each subregion over with and without sclerosis groups. The spot color represented an increase (red) or decrease (blue) of density in sclerosis. The cell types shown in (**b and c**) were marked in red. **(b)** The spatial distribution of subtypes of IN neurons with and without sclerosis. (**c**) The spatial distribution of VIP.0 and VIP.5 neurons with and without sclerosis. (**d**) Dotplot showing the expression of *APP* and *CD74* in the APP-CD74 interaction-related subtypes. (**e**) Dotplot showing the strength of the APP-CD74 interaction pairs with and without sclerosis. Red represents a strong interaction. (**f-g**) Heatmap displaying the expression of *VIP* and *VIP* receptors across cell types (**f**) and subtypes (**g**) with and without sclerosis.

**Extended Data Fig. 16.**
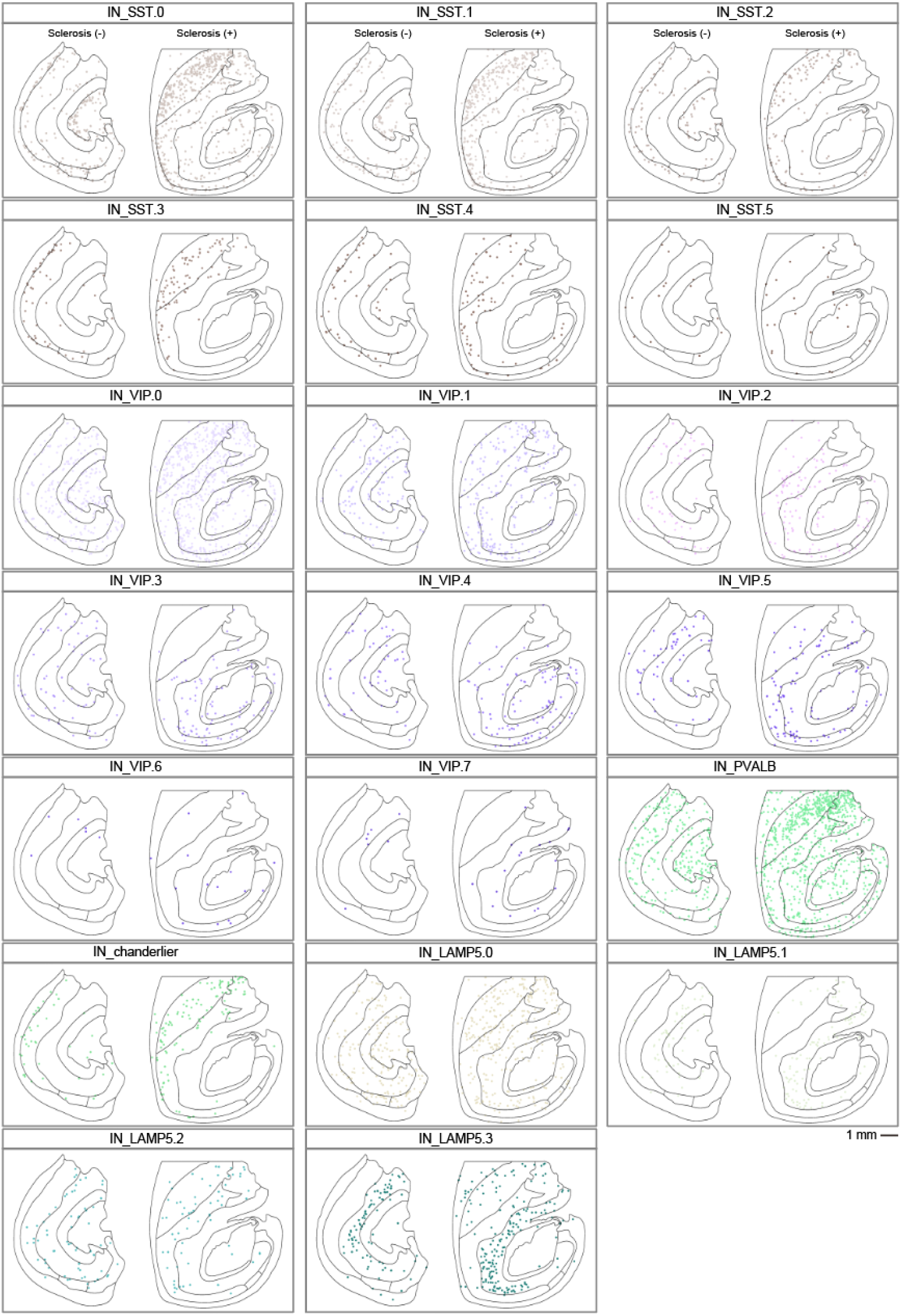
Spatial distribution of annotated subtypes of inhibitory neurons from Stereo-seq. Spatial visualization of each subtype of IN_SST, IN_VIP, IN_PVALB and IN_LAMP5.

